# Synergistic non-neutralizing plasma antibodies to PfRH5 drive potent malaria parasite growth inhibition

**DOI:** 10.64898/2025.12.24.696081

**Authors:** Jeffrey Marchioni, Kirsty McHugh, Jordan R. Barrett, Allison Seeger, Cassandra A. Rigby, Doris Quinkert, Ana Rodrigues, Anna Huhn, Dimitra Pipini, Angela M. Minassian, Jessica Kain, Douglas R. Townsend, Sean A. Knudson, Randall S. MacGill, George Georgiou, Simon J. Draper, Gregory C. Ippolito, Jason J. Lavinder

**Author notes:** Corresponding authors Contacts: J.J.L.; G.C.I.; S.J.D. Authors contributed equally (Jeffrey Marchioni, Kirsty McHugh).

## Abstract

An effective blood-stage vaccine is needed to protect against malaria pre-erythrocytic stage breakthrough. *P. falciparum* reticulocyte-binding protein homolog 5 (PfRH5) has emerged as a promising blood-stage vaccine antigen candidate, reducing parasite growth in humans during malaria challenge and showing field efficacy in children. Here, we characterize the human plasma IgG response to the RH5.1 vaccine candidate at monoclonal resolution, revealing that plasma repertoires are dominated by abundant, non-neutralizing antibodies. Using oligoclonal reconstitution experiments, in which defined pools of recombinant plasma mAbs are reassembled and functionally tested, we map how individual antibody interactions shape parasite growth inhibition activity. This approach allows us to discern which antibodies, within a polyclonal setting, act additively or synergistically, thereby revealing the emergent properties of anti-PfRH5 IgG. We further show that IgG lineages targeting linear epitopes lack neutralizing activity, while non-neutralizing IgG lineages that bind conformational epitopes can exhibit potent, interdependent synergy with each other and with neutralizing mAbs. These synergistic antibodies were identified in the plasma IgG compartments of five volunteers and highlight non-neutralizing PfRH5 epitopes that are critical for polyclonal-mediated growth inhibition. Our findings have broad implications for PfRH5 vaccine immunogen engineering and the role of non-neutralizing antibodies in infectious disease immunity.

## Introduction

Malaria is one of the deadliest diseases in the global south, resulting in an estimated 263 million infections and 597,000 deaths in 2023.^1^ Interventions such as netting, pesticides, and antimalarial drugs have alleviated or even eliminated malaria burden in many endemic countries, with 2.2 billion cases and 12.7 million deaths having been averted worldwide since 2000. However, the decline in malaria case incidence has been gradual since the 2010s, and the burden remains heavily apparent in sub-Saharan Africa, where in 2023 94% of global clinical malaria cases and 95% of global deaths were reported.^1^ Ultimately, the ongoing threat from increasing drug and pesticide resistance, as well as shifting malaria transmission patterns due to climate change, underscore that improved vaccines and prophylactics are critical to achieve sustained control of malaria.^2^

As of 2024, the World Health Organization recommends the programmatic use of two vaccines (RTS,S/AS01 and R21/Matrix-M) in young children in malaria endemic regions with moderate to high transmission. Both vaccines target the circumsporozoite protein (CSP) on sporozoites, which facilitate the pre-erythrocytic stage of malaria infection. R21/Matrix-M appears to have improved efficacy and longevity compared to RTS,S/AS01, providing hope that a low-cost vaccine can reduce malaria burden in the global south.^3,4^ Further, anti-CSP monoclonal antibodies (mAbs), like CIS43LS,^5,6^ L9LS,^7^ and MAM01,^8^ have shown safety and clinical success in preventing malaria infection. Although promising, pre-erythrocytic stage vaccines or mAbs typically do not provide long-term sterilizing immunity, allowing merozoites to enter the bloodstream and begin the cyclical blood-stage infection responsible for the most severe malaria pathology.^9^ To address this, blood-stage vaccines are in development to be assessed alone or in combination with current pre-erythrocytic stage candidates (i.e., multi-stage vaccination).

Previous blood-stage vaccine antigen candidates, such as merozoite surface protein-1 (MSP-1) or apical membrane antigen-1 (AMA-1), have failed in clinical trials to effectively prevent clinical malaria. Despite the discovery of neutralizing antibodies against these targets,^10,11^ they are limited by high levels of polymorphism,^12,13^ redundancy of red blood cell invasion pathways,^14^ and the requirement for high antibody concentrations to confer protection.^15^ More recently, the *P. falciparum* reticulocyte-binding protein homolog 5 (PfRH5) has emerged as a promising bloodstage vaccine antigen candidate.^16^ PfRH5 binds basigin (CD147) on the surface of erythrocytes and is known to form a pentameric invasion complex with the cysteine-rich protective antigen (PfCyRPA), RH5-interacting protein (PfRIPR), cysteine-rich, small, secreted (PfCSS), and *Plasmodium* thrombospondin-related apical merozoite protein (PfPTRAMP), together called the PCRCR-complex.^17,18^ All members of this complex are essential for merozoite invasion in *P. falciparum*^17,19^ and PfRH5, PfCyRPA, and PfRIPR are known to have low polymorphism frequencies.^20^ Neutralizing antibodies and nanobodies have been found that target all five members of the complex,^17,19,21–24^ indicating all as potential antigen targets for blood-stage immunity.

Within the PCRCR-complex, PfRH5 has been studied extensively as a vaccine antigen candidate, and notably, formulating RH5.1 protein in AS01_B_ adjuvant increased anti-PfRH5 serum IgG concentrations in vaccinated malaria-naïve United Kingdom (UK) adult volunteers ∼10-fold as compared to original trials using a viral-vectored delivery platform.^25–27^ Upon challenge via controlled human malaria infection (CHMI), UK adults vaccinated with RH5.1/AS01_B_ exhibited a significant reduction in parasite growth, a milestone for the blood-stage vaccine field.^27^ Subsequently, the same RH5.1 protein vaccine reformulated in Matrix-M adjuvant showed partial efficacy against clinical malaria in 5-17 month old African children in an ongoing Phase 2b trial.^28^ In a follow-up study, >200 mAbs isolated from B cells of UK adult vaccinees were characterized.^29^ All neutralizing mAbs targeted conformational epitopes on PfRH5 proximal to or directly competing with the PfRH5-basigin binding site. Of the non-neutralizing mAbs, a subset of them exhibited synergistic properties that enhanced the potency of the neutralizing mAbs in a growth inhibition activity (GIA) assay. This effect is thought to result from these non-neutralizing mAbs slowing merozoite invasion into erythrocytes, thereby providing more time for the neutralizing antibodies to bind PfRH5, as previously described.^30^

Although the properties of many neutralizing and non-neutralizing mAbs derived from PfRH5-specific B cells have been well characterized, their abundances and functionality within the vaccine-induced pool of circulating IgG remain poorly defined. Using Ig-Seq,^31–33^ we applied high-resolution bottom-up proteomics to delineate the polyclonal plasma anti-PfRH5 IgG response of malaria-naïve UK adults following RH5.1/AS01_B_ vaccination. We profiled these circulating antibodies in terms of abundance, specificity, and functional properties. Notably, the plasma IgG response to PfRH5 was dominated by non-neutralizing antibody lineages, several of which acted synergistically with both neutralizing and non-neutralizing antibodies to significantly enhance blood-stage parasite growth inhibition.

## Results

### Proteomic delineation of circulating anti-PfRH5 IgG after RH5.1 vaccination

Malaria-naïve UK adult volunteers (n=5) were administered three monthly doses of 10 µg RH5.1 protein formulated in 0.5 mL AS01_B_ adjuvant, eliciting a mean anti-PfRH5 plasma IgG concentration of 114.5 µg/mL at day 69 (13 days post-3rd immunization).^27^ On day 69 samples, we employed high-resolution bottom-up tandem mass spectrometry (Ig-Seq) to characterize the RH5.1-specific plasma-derived IgG response to vaccination.^31,32^ In parallel, we also performed Illumina MiSeq next-generation sequencing of both natively-paired (VH:VL)^34^ and separate antibody heavy (VH) and light chain (VL) variable regions derived from day 63 peripheral B cells (BCR-Seq)^32^ (Fig. 1A). These combined approaches enabled the identification, relative quantitation, recombinant expression, and functional characterization of antigen-specific plasma antibodies. Plasma IgG lineages, clustered by CDRH3 homology, are quantified in terms of relative abundance of the PfRH5-specific IgG response, determined from extracted ion chromatogram (XIC) precursor areas of CDRH3 peptides within each lineage (Fig. 1B and Fig. S1). The plasma antibody repertoire was largely oligoclonal across all five donors, with the top 50% (D50) of PfRH5-reactive IgG (as determined by Ig-Seq) comprising a mean of ∼9 lineages. PfRH5-reactive plasma IgG lineages demonstrated mean somatic hypermutation of 2.9% and mean CDRH3 length distributions of 16 amino acids (Fig. S2), consistent with previously examined PfRH5-specific B cell responses in these donors.^29^

**Figure 1.**
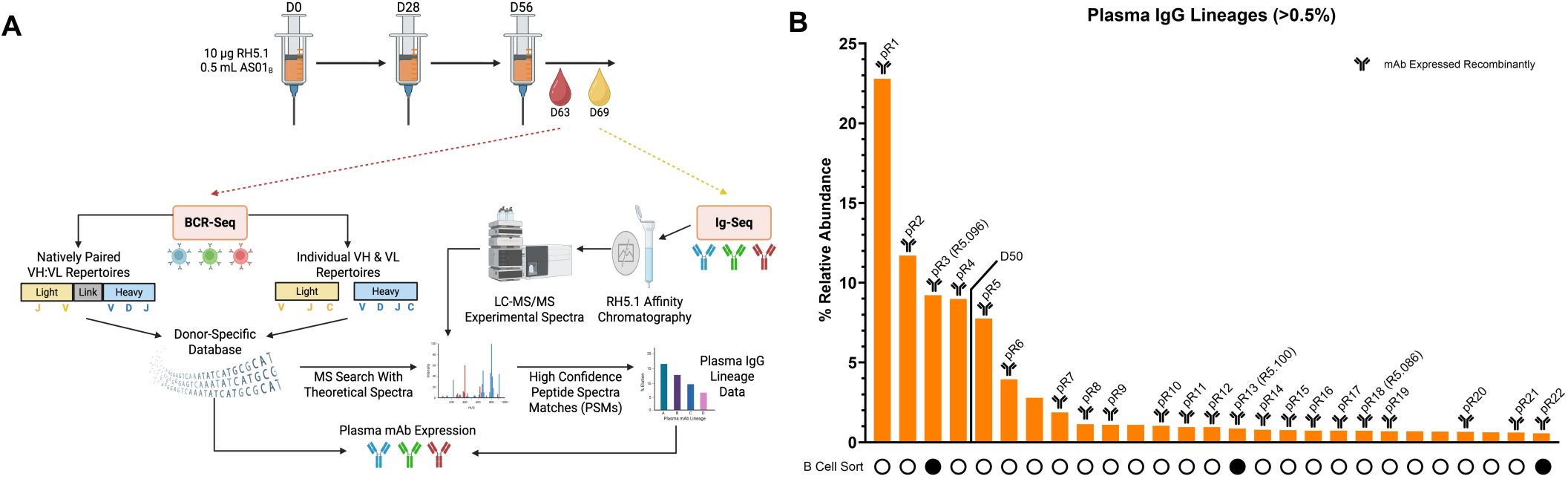
The Anti-RH5.1 Plasma IgG Repertoire. (A) Timing of immunizations and sample collections for BCR-Seq and Ig-Seq analyses. Created in BioRender. Marchioni, J. (2025) https://BioRender.com/j0rmh3w. (B) The relative abundances of RH5.1-specific IgG lineages in Donor A. The number of lineages that account for ∼50% of the anti-RH5.1 plasma IgG repertoire is denoted by a line labeled with “D50.” Below each bar, a filled in circle acknowledges that the lineage CDRH3 has been identified prior using RH5-specific B-cell sorting.^29^

For one of the donors (Donor A), we identified the paired VH and VL sequences of 22 PfRH5-specific plasma IgG lineages present at ≥0.5% relative abundance (Fig. 1B). These 22 lineages represented ∼79% relative abundance of the total PfRH5-reactive IgG detected, thereby enabling recombinant expression of the majority of the vaccine-induced plasma antibody response in this donor. Further, for the four other donors (Donors B-E), we recombinantly expressed an additional 13 highly abundant plasma antibody lineages (Data S1 and Fig. S1), including the top-ranked plasma IgG in each donor (8.9 – 19.8% relative abundance). In total, 34 of the 35 recombinantly expressed plasma mAbs bound to the RH5.1 target by indirect ELISA (pR5N1 did not bind). Of note, in a separate study on this donor cohort, Barrett *et al.* previously sampled four of these plasma IgG lineages (pR3, pR13, pR18, and pR33) as recombinant mAbs from antigen-specific B cells (R5.096, R5.100, R5.086, and R5.239, respectively, in reference^29^).

### Epitope mapping and functional profiling reveals dominance of non-neutralizing plasma IgG

Using high-throughput surface plasmon resonance (HT-SPR), we next measured the binding kinetics of all 34 recombinant plasma IgG mAbs against RH5.1 (Data S1). Plasma IgG were consistently high affinity, with *K*_D_ values ranging between ∼60 pM and 3 nM, with a mean value of 810 pM (Fig. S3). Additionally, anti-PfRH5 plasma mAbs, along with a set of previously characterized control mAbs from antigen-sorted B cells,^29,30^ were analyzed in a HT-SPR pairwise competitive binding assay to group them into epitope bins (Fig. S4). This enabled the PfRH5-specific plasma IgG response to be visualized as a community network plot, illustrating individual mAb communities and their competitive binding interactions (Fig. 2A).

**Figure 2.**
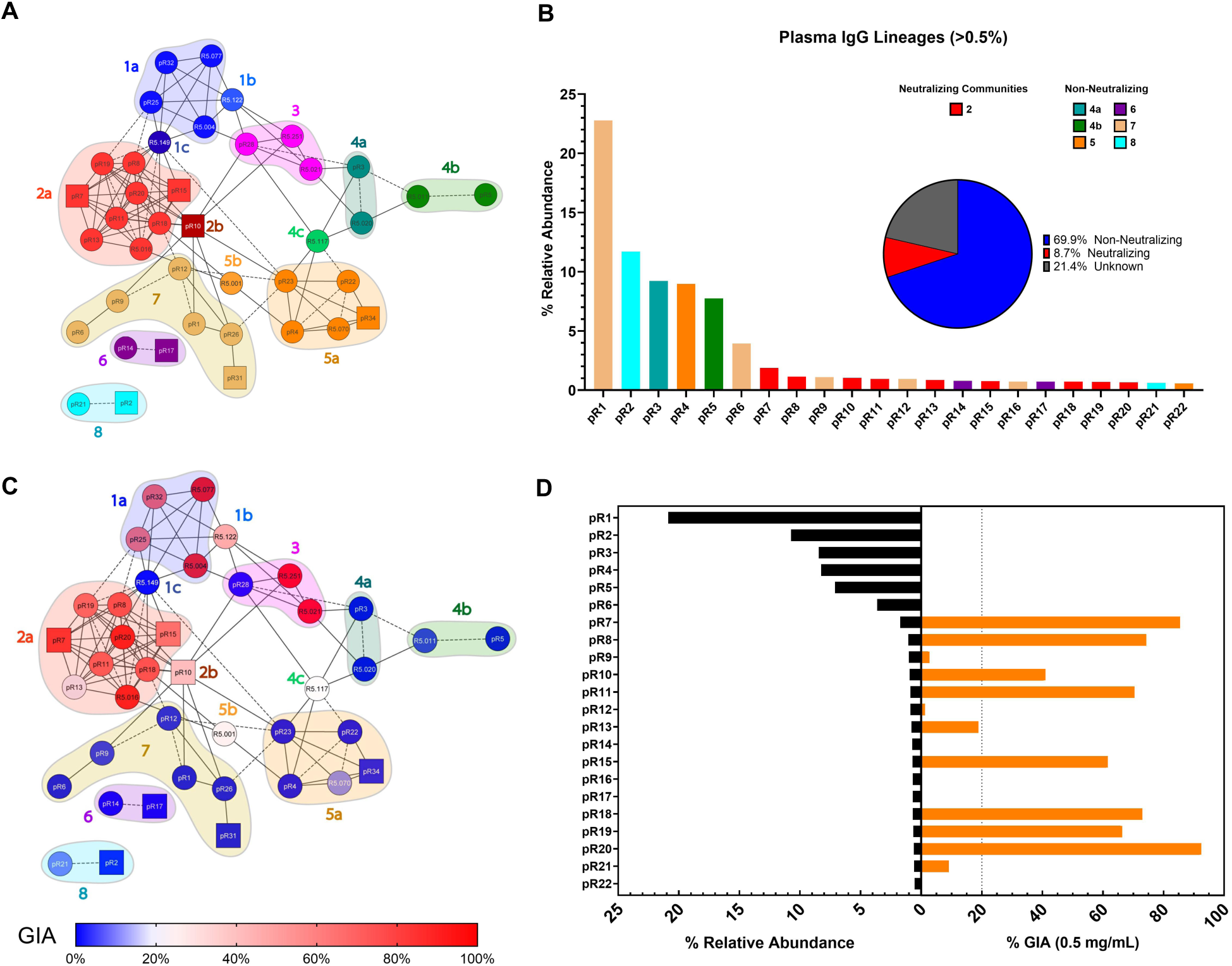
Functional Characterization of anti-RH5.1 Plasma mAbs. (A) Community network plot identifying pairwise competition among plasma mAbs, determined using high-throughput surface plasmon resonance (HT-SPR). Each community is identified by a color and number code, and subcommunities indicated by a letter. Individual mAbs are represented by nodes. Control mAbs^29,30^ are listed with the prefix “R5,” while plasma mAbs are listed with the prefix “pR.” Dashed lines and solid lines indicate unidirectional and bidirectional competition, respectively. Square nodes indicate that a mAb was excluded as either ligand or analyte. Several mAbs were excluded from analysis due to poor binding kinetics, yet some epitopes were resolved using PfRH5 variant ELISAs (Fig. S5). (B) Epitope binning of plasma mAbs from Donor A (Fig. 1B) in relation to the relative abundance of their respective lineages. mAbs are color-coded by epitope communities from (A). The pie chart shows the relative plasma abundance of neutralizing/GIA-positive (communities 1-3) or non-neutralizing/GIA-negative (communities 4-8) mAbs found at ≥0.5% relative abundance. Lineages not expressed as mAbs, due to lack of VH:VL information or low relative abundance, are denoted in gray as unknown. (C) mAb percent GIA, at 0.5 mg/mL, superimposed upon community network plot in (A). A value ≥20% (red) indicates GIA-positive mAbs, while those below (blue) indicate GIA-negative mAbs. (D) The relative abundance of Donor A’s mAbs from (B) compared to percent GIA of individual mAbs tested at 0.5 mg/mL from (C).

The plasma IgG response to PfRH5 consisted of lineages spanning six major epitope bins on the PfRH5 protein, which had been previously defined.^29^ Antibodies within epitope communities 1-3 surround the “diamond tip” of PfRH5, overlapping or proximal to the PfRH5:basigin binding site, and are often neutralizing. mAbs targeting non-neutralizing epitopes bind distal to the “diamond tip” and include epitopes targeted by synergistic mAbs (community 4), the PfRH5:PfCyRPA binding site (community 5), and the intrinsic disordered loop (IDL, community 6) of PfRH5. Relative to prior binning results,^29^ the plasma IgG response resolved an additional PfRH5 sub-community, designated 2b, which exhibits a distinct, though related, binding profile compared to all other community 2 antibodies (now designated as 2a). Additionally, due to a smaller population of mAbs experimentally binned, community 3 did not resolve into separate sub-communities 3a and 3b. Further, we identified plasma IgG targeting two additional linear epitope regions, confirmed by indirect ELISA to be the disordered N- and C-termini of PfRH5, defining communities 7 and 8, respectively (Fig. S5). To the best of our knowledge, no N-terminal and only one C-terminal-specific mAb, R5.CT1,^35^ have been reported to date. Of note, 3 of 7 N-terminus and 2 of 2 C-terminus plasma IgG lineages utilize the IGHV5-51 gene, which was the most frequently utilized germline IGHV in the PfRH5-specific plasma IgG response across all five donors examined (Data S1 and Fig. S6).

Next, we examined the relationship between binning and circulating IgG quantification data to detail the epitope specificity of the most abundant PfRH5-specific plasma mAbs in Donor A (Fig. 2B). The six most abundant mAbs (pR1-6), which together account for ∼64% relative abundance (3.9 – 22.8% individually) of the plasma IgG response in this donor, binned within non-neutralizing epitope communities 4-8. Notably, although prior studies had reported that neutralizing (GIA-positive) community 1 and 3 antibodies can be readily identified within the circulating the B cell repertoire,^29^ only neutralizing mAbs from community 2 were identified within this donor. Individually, community 2 mAbs were relatively low abundance, all being detected at <2% of the total PfRH5-specific plasma IgG response, whereas combined, they accounted for ∼8.7% of the response. Overall, plasma mAbs targeting non-neutralizing epitopes were found to circulate at ∼8x greater concentration than those targeting neutralizing epitopes (Fig. 2B inset).

To expand our analysis, we mapped the GIA of plasma IgG mAbs (Data S1) across all five donors onto the PfRH5 epitope network plot (Fig. 2C). Plasma IgG antibodies in communities 1a and 2 were, as expected, found to be neutralizing, whereas the lone plasma mAb assigned to community 3 (pR28) was non-neutralizing (i.e., had no GIA). Plasma IgG antibodies that binned to communities 4-6, or those that target the N- or C-terminal regions (communities 7 and 8, respectively), were all GIA-negative (<20% GIA). Of note, several plasma mAbs from community 1 (pR25 and pR32), community 2 (pR7 and pR11), and community 7 (pR1, pR12, and pR31) blocked the PfRH5:basigin interaction (Fig. S7). Interestingly, although some community 7 (N-terminus) mAbs demonstrated basigin blocking, they did not demonstrate GIA. Four community 5 mAbs (pR4, pR22, pR23, and pR34) blocked the PfRH5:CyRPA interaction and did not have any GIA responses (Fig. 2C and S7), consistent with previous studies.^29,36^ Within Donor A exclusively (Fig. 2D), we confirmed that the top six plasma mAbs are non-neutralizing when tested individually and that all eight GIA-positive (≥20% GIA) plasma IgG mAbs belong to community 2.

Across four of the five donors, the most abundant plasma IgG mAbs (pR1, pR23, pR26, and pR29) were all non-neutralizing (GIA-negative) (Fig. 2C and S8). In the remaining donor, the third most abundant IgG mAb (pR33, 10.2% relative abundance), targeted the IDL of community 6 and was also GIA-negative (Fig. S8). Altogether, these findings reveal that the anti-PfRH5 plasma IgG response across multiple donors is dominated by non-neutralizing antibodies.

### Abundant non-neutralizing plasma IgGs play an essential role in polyclonal potency

To investigate whether the most abundant plasma IgG, while non-neutralizing, may contribute to overall polyclonal neutralization potency, we designed GIA oligoclonal reconstitution experiments (OREs) that contained subsets of the PfRH5-specific plasma neutralizing and non-neutralizing mAbs at their respective plasma concentrations (Ig-Seq abundances) (Fig. 3A and S9). Pool 1 (GIA EC_50_ = 113.7 µg/mL; Fig. 3B) corresponds to the complete pool of recombinant PfRH5-specific plasma mAbs, whereas in pools 2-13, one or more of the mAbs from pool 1 was replaced with a “decoy” negative-control IgG to maintain the same total IgG concentration across all OREs. GIA titration curves from OREs were used to calculate EC_50_ and EC_80_ values to statistically compare pools with each other (Fig. 3B and 3C).

**Figure 3.**
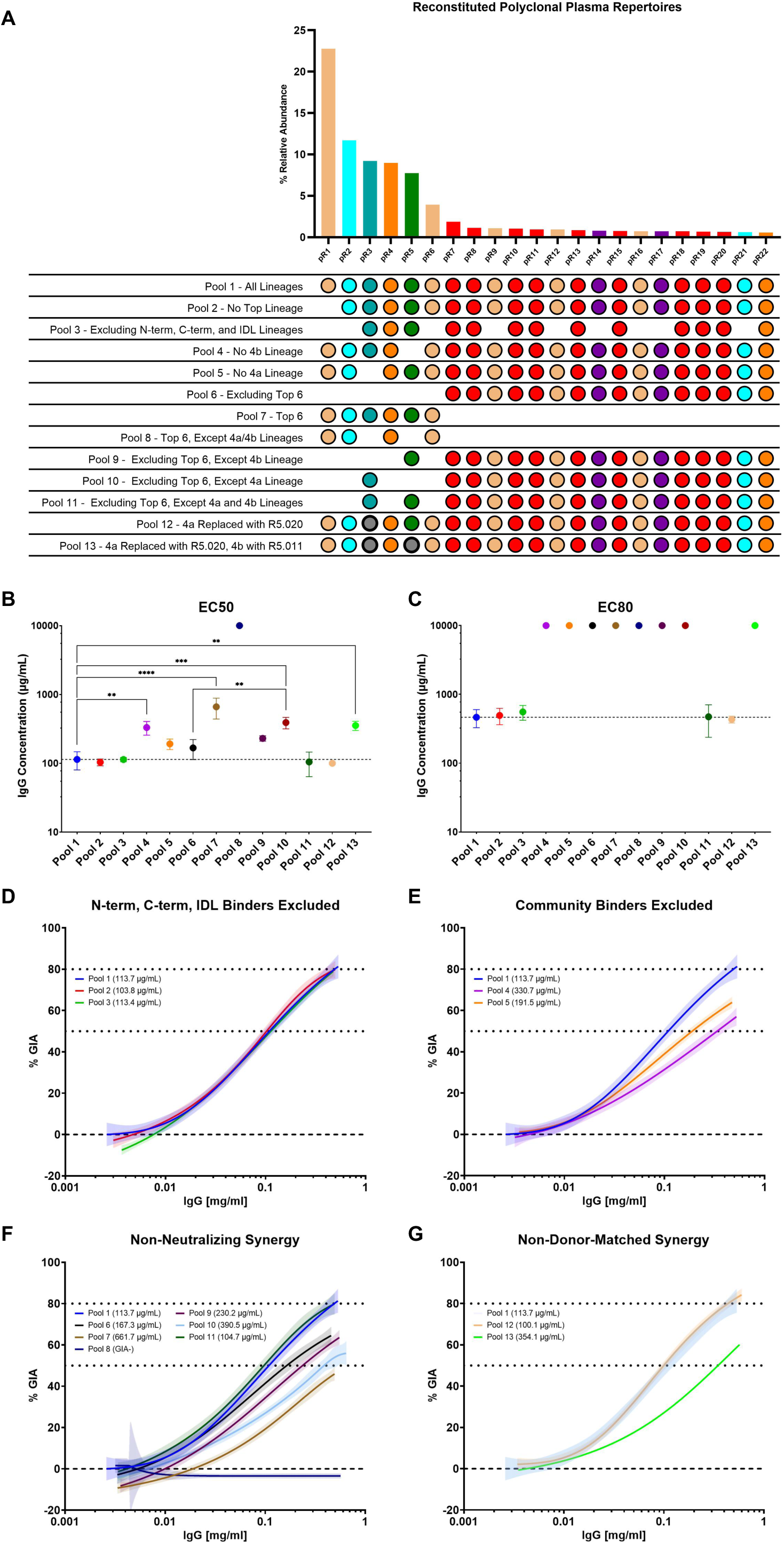
Oligoclonal Reconstitution Experiments (OREs) of Plasma IgG mAbs. (A) Schematic describing the composition of reconstituted pools used in GIAs. A missing bubble indicates a mAb was excluded. Pools were mixed to a final concentration of 0.5 mg/mL and used in 8-step, 2-fold serial dilution GIA assays. Individual mAbs were diluted to concentrations based on normalized relative plasma proteomic abundance (determined by Ig-Seq). Excluded mAbs were replaced with a negative control “decoy” mAb at the concentration needed to reach a pool concentration of 0.5 mg/mL. (B) EC_50_ and (C) EC_80_ values for all pools were interpolated from the dilution curves of triplicate experiments (Fig. S8), each using erythrocytes from distinct blood donor volunteers. (B-C) The dashed line indicates the EC_50_ or EC_80_ value of pool 1. Weak or non-active mAb pools, for which EC values could not be determined, were assigned values of 10 mg/mL for the purpose of analysis. * P < 0.05, ** P < 0.01, *** P < 0.001; **** P < 0.0001, determined by ANOVA with Tukey’s multiple comparison post-test. Growth inhibition curves of three triplicate experiments representing ORE pool 1 with (D) pools 2-3, (E) pools 4-5, (F) pools 6-11, and (G) 12-13. EC_50_ values are listed in parentheses next to each pool in the legend. (D-G) All data points from three independent experiments were combined and fit using a single non-linear regression model (pooled fit) to visualize the overall trend and 95% confidence intervals are plotted.

The most abundant plasma mAb from pool 1 (pR1, 22.8% relative abundance) binds to the PfRH5 N-terminus (community 7) and has no GIA activity on its own. As expected, removing it from pool 1 had no significant impact on GIA (Fig. 3D, pool 2; EC_50_ = 103.8 µg/mL). Plasma mAbs that bind to linear, disordered epitopes on PfRH5, such as the N-terminus, C-terminus, or IDL, namely communities 6-8, collectively comprise 43.3% of anti-PfRH5 plasma IgG abundance (Fig. 2B) and are non-neutralizing when tested individually by GIA (Fig. 2D). Accordingly, removing all mAbs that bind these epitopes resulted in GIA essentially identical to that observed with pool 1 (Fig. 3D, pool 3; EC_50_ = 113.4 µg/mL). Collectively, these results reveal that IgG targeting linear, disordered epitopes on PfRH5 are non-neutralizing in both a monoclonal and polyclonal setting and do not contribute toward overall polyclonal-mediated GIA via allosteric or other indirect effects.

The plasma IgG mAbs pR3 and pR5 map to non-neutralizing epitope communities 4a and 4b, respectively, and prior studies indicated that community 4 mAbs often have synergistic or antagonistic properties when tested in combination with neutralizing and non-neutralizing mAbs. For example, community 4a mAbs have been shown to antagonize the GIA of community 2 and 3b mAbs and synergize with community 1 and 3a mAbs. On the other hand, community 4b mAbs synergize primarily with community 1 and 3 mAbs.^29^ To evaluate the contribution of these community 4 mAbs within this donor’s IgG repertoire, we tested their individual effects in OREs (Fig. 3A and 3E). Removing pR5 (community 4b) from pool 1 resulted in a significant, ∼3-fold decrease in GIA potency (Fig. 3E, pool 4; EC_50_ = 330.7 µg/mL; ** p < 0.01). This finding was unexpected because, as mentioned above, earlier studies had reported that community 4b antibodies primarily synergize with neutralizing antibodies in communities 1 and 3, rather than those found in community 2 (the only neutralizing mAbs in pool 1). Similarly, when we removed pR3 (community 4a) from pool 1, there was a reproducible, yet statistically non-significant, decrease in neutralization potency (Fig. 3E, pool 5; EC_50_ = 191.5 µg/mL). However, the synergistic contribution of these plasma mAbs was much more pronounced when considering the EC_80_ values (Fig. 3C), as neither pool reached the original maximal GIA potency of pool 1 by the top concentration tested. These findings reveal that both non-neutralizing community 4a and 4b plasma mAbs are contributing towards overall polyclonal GIA through synergistic interactions.

We next prepared a series of pools to further investigate the role of these non-neutralizing, synergistic plasma IgG mAbs. We first removed from pool 1 the six topmost abundant mAbs (pR1–6, together ∼64.4% relative abundance), which are all non-neutralizing individually (Fig. 2D and Fig. 3A). This resulted in a slight decrease in GIA potency that failed to reach statistical significance (Fig. 3F, pool 6; EC_50_ = 167.3 µg/mL). Unexpectedly, a pool containing only pR1-6 exhibited low, but dose-dependent GIA (Fig. 3F, pool 7; EC_50_ = 661.7 µg/mL), indicating synergy is occurring amongst mAbs in this pool even in the absence of individually neutralizing antibodies. To confirm our hypothesis that this was driven by the community 4 mAbs within this pool, both pR3 and pR5 were dropped from pool 7, leaving only pR1, pR2, pR4, and pR6 (Fig. 3F, pool 8), which had no detectable GIA. Combinations of pR3 and pR5 with each other, as well as with pR1, pR2, and pR4 confirmed that non-neutralizing community 4 mAbs pR3 and pR5 synergize with each other and no additional synergy occurs among the remaining non-neutralizing mAbs (Fig. S10). Next, each community 4 mAb was individually reintroduced into pool 6, which lacks the six topmost abundant IgG. Adding community 4b pR5 back into pool 6 resulted in no significant change in GIA (Fig. 3F, pool 9; EC_50_ = 230.2 µg/mL), indicating that pR5 alone is not sufficient to restore the GIA seen in pool 1. Surprisingly, separately reintroducing community 4a pR3 back into pool 6 resulted in a significant decrease in GIA potency (Fig. 3F, pool 10; EC_50_ = 390.5 µg/mL; p < 0.01). Even more remarkable, when both pR3 and pR5 were simultaneously returned to pool 6, the original GIA potency of pool 1 was fully restored, with no significant differences observed (Fig. 3F, pool 11; EC_50_ = 104.7 µg/mL). This suggests an interdependent, synergistic relationship between the non-neutralizing 4a and 4b mAbs, pR3 and pR5, alongside the neutralizing mAbs present at lower abundances.

Finally, to determine whether the synergism resulting from pR3 and pR5 is dependent upon properties unique to these particular mAb clones, we substituted pR3 and pR5 with non-donor matched community 4a and 4b mAbs previously isolated from sorted B cells, R5.020^29^ and R5.011,^30^ respectively. Our previous HT-SPR data confirmed that community 4a R5.020 competes with pR3 but not with pR5 (Fig. 2A) and replacing pR3 with R5.020 resulted in no significant difference in GIA as compared to pool 1 (Fig. 3G, pool 12; EC_50_ = 100.1 µg/mL). Prior characterization of mAbs from sorted B cells found that all community 4b mAbs tested compete with one or multiple community 4a mAbs.^29^ Community 4b R5.011 follows this trend by competing with both pR5 and pR3. Replacing both pR3 and pR5 with R5.020 and R5.011, respectively, resulted in a significant decrease in potency compared to Pool 1 (Fig. 3G, pool 13; EC_50_ = 354.1 µg/mL; p < 0.01). The ability of pool 12, but not pool 13, to sustain the polyclonal GIA of the completely reconstituted pool 1 highlights that non-competition between community 4 mAbs is required to elicit non-neutralizing GIA like that observed between pR3 and pR5. In summary, only when both community 4a pR3 and 4b pR5 mAbs were present (pools 2, 3, 10), or a similar mAb was substituted (pool 12), were we able to capture sufficient neutralization (≥80% GIA) to replicate the potency witnessed in pool 1 (Fig. 3C).

### Non-neutralizing community 4 mAbs synergize GIA potency of both donor- and non-donor-matched antibodies

Given their prevalence within the polyclonal antibody response, alongside their individual GIA-positive activity (Fig. 2B and Fig. 2D), each donor-matched community 2 mAb represented in the ORE pools (Fig. 3) was mixed with either pR3, pR5, or both in combination to delineate pairwise synergistic (or antagonistic) interactions within the polyclonal plasma response (Fig. S11A). To define synergism or antagonism, predicted Bliss additivity (% GIA) was calculated for each combination of mAbs tested. The synergy index, defined as the Bliss additivity value subtracted from the averaged measured % GIA of the mAb combination, identified whether a mAb combination was additive (index ∼ 0), synergistic (index ≥ 10%) or antagonistic (index ≤ −10%) (Fig. 4A). The donor-match community 2 mAbs fell into two groups. The first group (pR7, pR8, pR11, and pR18) demonstrated only additive GIA effects when combined with pR3 and pR5, and, in some cases, were antagonized by either pR3 or pR5 alone (Fig. 4A). The second group (pR10, pR13, pR19, and pR20) were synergized by pR3 and pR5 in combination. These results reinforce the ORE observations that pR3 and pR5, in combination, strongly synergize with many neutralizing community 2 mAbs within the plasma, positioning the non-neutralizing pair of community 4 mAbs as an essential driver of polyclonal GIA potency in this donor.

**Figure 4.**
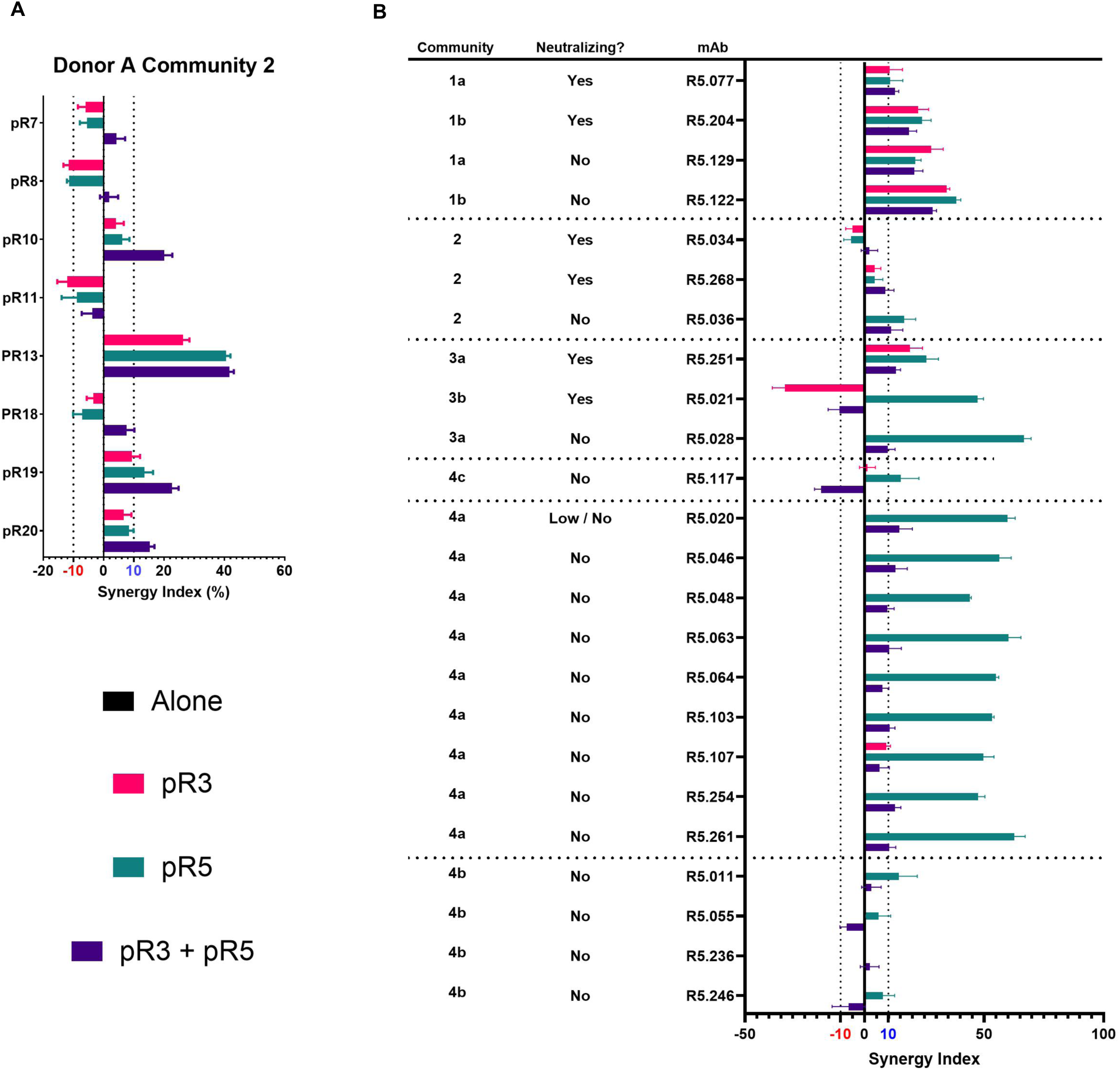
Plasma mAb Synergy and Antagonism of Neutralizing and Non-Neutralizing mAbs. (A) Synergy index (%) of community 2 mAbs used in OREs in Figure 3 with either Community 4a pR3, Community 4b pR5, or both (0.4 mg/mL combined). (B) Synergy index of previously defined neutralizing and non-neutralizing mAbs^29,30^ with either Community 4a pR3, Community 4b pR5, or both. Dashed lines indicate grouping of separate epitope communities. (A-B) The synergy index is calculated as the predicted Bliss additivity (% GIA) subtracted from the average measured % GIA of the mAb combinations. This value defines the combination of mAbs as synergistic (≥ 10%), antagonistic (≤ −10%), or additive, with dashed lines drawn to delineate thresholds.

To assess whether these observed synergistic effects were unique to donor-matched, plasma-derived mAbs (i.e., a potential donor-specific polyclonal effect), we performed pairwise synergy screens with previously characterized B cell-derived mAbs from other donors.^29,30^ GIA potency and synergy indices were calculated for each pair (Fig. 4B and S11B). When tested independently, both pR3 and pR5 exhibited synergy with the neutralizing mAbs tested from other donors in epitope communities 1a, 1b, and 3a, and non-neutralizing mAbs in communities 1a and 1b, patterns consistent with previously documented community 4a and 4b mAbs.^29^ These trends were maintained when pR3 and pR5 were evaluated in combination. Distinct behaviors emerged for other communities. The community 3b mAb R5.021 was antagonized (or likely competed with, Fig. 2A) by pR3 but strongly synergized by pR5. Among the three tested community 2 mAbs, only non-neutralizing R5.036 GIA was synergized, by pR5 alone and the pR3 and pR5 combination. Importantly, pR3 did not reproduce its synergistic behavior with pR5 when tested with four non-donor matched community 4b mAbs (R5.011, R5.055, R5.236, and R5.246). In stark contrast, pR5 synergized with all nine community 4a mAbs examined, distinguishing it from previously described 4b mAbs. Together, these data underscore the unique functional contributions of non-neutralizing, synergistic community 4 mAbs within the polyclonal PfRH5-specific plasma IgG responses.

### RH5.1/AS01_B_ immunization elicits synergistic community 4 plasma IgG lineages in all five donors

To evaluate whether community 4 synergy is a recurring feature in the RH5.1/AS01_B_ vaccine-induced PfRH5-specific plasma IgG repertoire, we attempted to identify additional 4a and 4b lineages across the remaining four donors’ Ig-Seq data. To select sequences, we searched our database for lineages that share previously described VH and VL usage^29,30^ for these epitope communities, in which community 4 mAbs utilize predominantly IGHV3 (mostly IGHV3-7 and IGHV3-21) or IGHV2-70 and IGLV3-21. One of these plasma-derived mAbs, pR50, was previously identified using B cell sorting (R5.048)^29^ and is already known to compete with community 4a. An additional 12 mAbs detected in the PrRH5-specific plasma IgG repertoire of Donors B-E were selected that utilize these shared, public community 4 germline features (Fig. 5A). To validate our selection prior to recombinant expression, we utilized AlphaFold3^37^ to predict the structural interaction of each mAb with RH5ΔNL (RH5 lacking the disordered N-terminus and IDL). Predictions for mAbs pR51, pR52, pR60, and pR61 demonstrated binding to the community 4a epitope, yielding ipTM scores ranging from 0.77 to 0.8 (Fig. S12), while all other mAbs had poor prediction profiles.

**Figure 5.**
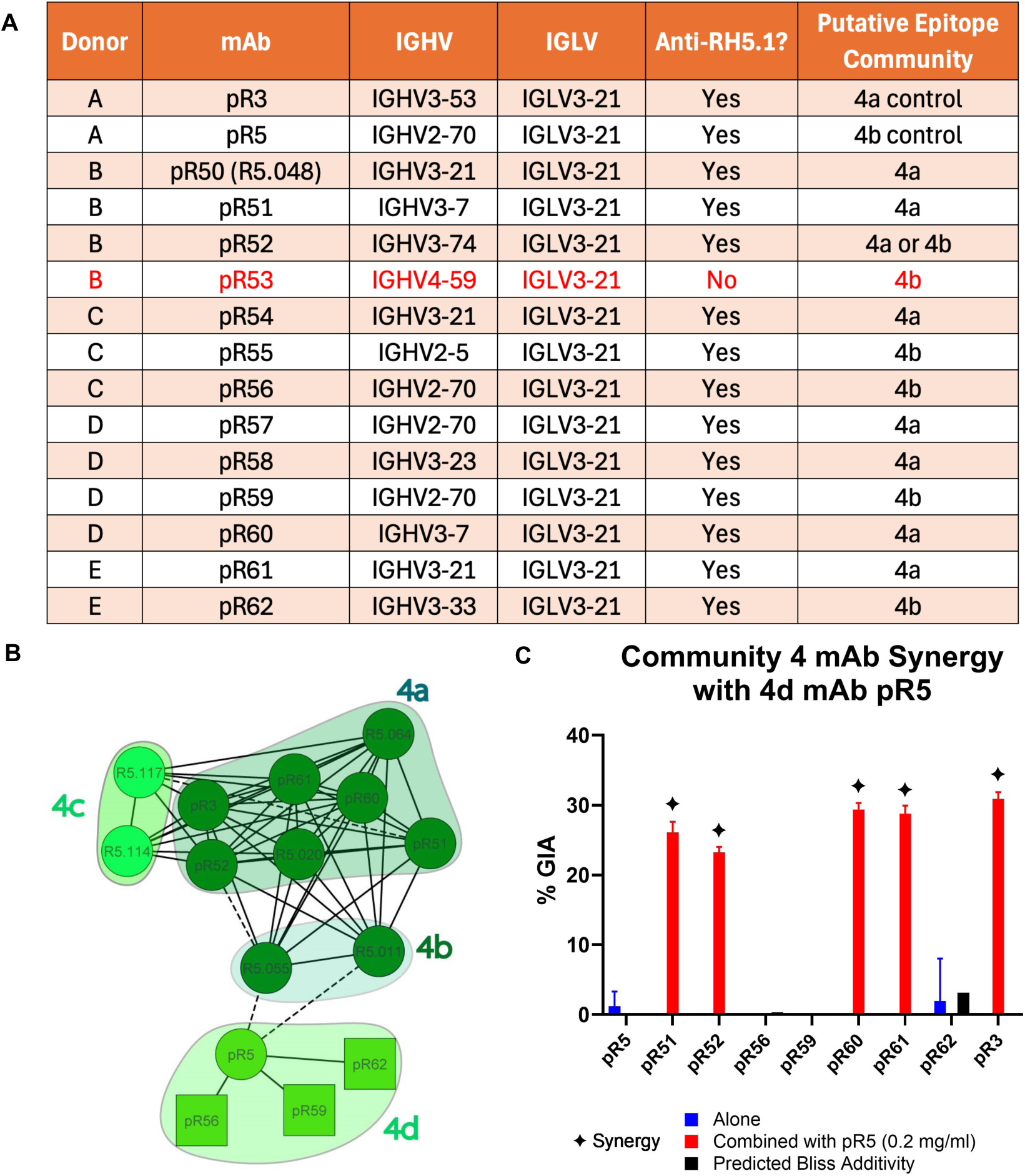
**Community 4a/4b Synergy Screens**. (A) Putative community 4a or 4b mAbs pR50-62 were selected based on previously discovered IGHV and IGLV gene combinations and tested for reactivity against RH5.1 via ELISA. pR53, in red, did not bind to RH5.1 protein via ELISA. (B) Community network plot identifying pairwise competition among putative community 4a, 4b, and 4d plasma mAbs, determined using HT-SPR. Individual mAbs are represented by nodes. Control mAbs^29,30^ are listed with the prefix “R5,” while plasma mAbs are listed with the prefix “pR.” Dashed lines and solid lines indicate unidirectional and bidirectional competition, respectively. Square nodes indicate that a mAb was excluded as either ligand or analyte. For full network plot, see Fig. S13. (C) pR51-pR61 were tested for GIA synergy with community 4d mAb pR5. Test mAbs (including positive control pR3) were held at 0.5 mg/mL while pR5 was held at 0.2 mg/mL. Blue bars represent the GIA value of the test mAb alone, red bars represent the test mAb combined with pR5, and black bars represent predicted Bliss additivity (% GIA) of the test mAb combined with pR5. The blue bar at the far left of the plot indicates the GIA of pR5 alone. The synergy index is calculated as the predicted Bliss additivity (% GIA) subtracted from the average measured % GIA of the mAb combinations. The combination of the two mAbs were defined as synergistic if GIA ≥ 10%, (filled-in star).

Upon expression, 11 of 12 mAbs bound to RH5.1 by indirect ELISA. We next examined 11 putative 4a/4b mAbs (excluding pR5.050) via a competitive indirect ELISA, with biotinylated pR3 and pR5 as competition controls (Fig. S13). Four mAbs (pR51, pR52, pR60 and pR61) competed exclusively with pR3, while two (pR59 and pR62) competed exclusively with pR5. pR56 competed with both pR3 and pR5, showing a stronger competition index for pR5. These mAbs were further analyzed by HT-SPR in pairwise competitive binding assay with pR1-pR34 and previously characterized control mAbs from antigen-sorted B cells (Fig. S14).^29,30^ Consistent with the competition ELISA (Fig. S13), pR51, pR52, pR60, and pR61 binned with community 4a mAb pR3. Unexpectedly, pR5, pR56, pR59, and pR62 now binned separately from the community 4b mAb R5.011 (Fig. 5B); we therefore designate these mAbs as belonging to a new subcommunity, community 4d. The remaining mAbs (pR54, pR55, pR57, and pR58), fell in other communities, including communities 2 and 5 (Fig. S14). This confirmed that community 4 lineages can be readily identified as plasma IgG across all donors, with both 4a and 4d (previously 4b) subcommunities found in Donor A (pR3 and pR5), Donor D (pR60 and pR59), and Donor E (pR62 and pR61). Additionally, positive identification of mAbs within both communities reinforce evidence that public genes exist for both communities 4a and 4b/4d, consistent with prior mAbs identified.^29^

The functional properties of these mAbs were further validated in a synergy GIA assay, where community 4a mAbs were tested with pR5 (Fig. 5C). Consistent with the epitope binning results (Fig. 5B), we found that pR51, pR52, pR60, and pR62 are individually non-neutralizing, but exhibit synergy with pR5 to exhibit GIA-positive behavior. All together, these findings highlight the potential for synergistic plasma antibody interactions to have profound effects on GIA, induced by the polyclonal antibody response to RH5.1 vaccination.

## Discussion

PfRH5 has emerged as a highly promising blood-stage antigen target for vaccine development,^25,27,28,35,38–40^ becoming the first member of the PCRCR-complex to enter clinical trials. In a Phase 1/2a clinical trial, RH5.1/AS01_B_ has shown pivotal evidence that PfRH5 immunization reduces parasite multiplication rates (PMR) following a controlled human malaria infection challenge in malaria-naïve UK adults.^27^ In a subsequent Phase 2b clinical trial with RH5.1/Matrix-M, protective efficacy up to 55% against clinical malaria was seen in Burkinabe children over a 6 month malaria transmission season. Both trials represent critical milestones in the blood-stage malaria vaccine field and highlight the promise of anti-PfRH5 immunity against *P. falciparum* in endemic regions. Despite these advancements, further improvements in neutralizing anti-PfRH5 antibody responses are required for higher-level protection,^27,41^ highlighting the need for next-generation vaccine immunogens targeting PfRH5 or its associated complex. Achieving this goal necessitates a detailed understanding of the RH5.1-elicited plasma antibody repertoire regarding its composition, dominant lineages, targeted epitopes, and the functional interactions that shape the polyclonal response.

While previous studies have focused on monoclonal antibodies derived from peripheral B cells and the structured core of PfRH5 (RH5ΔNC),^29^ our investigation surveys the circulating plasma IgG repertoire specific to the intact full-length RH5.1 immunogen. This distinction is important for two reasons: 1) identification and characterization of PfRH5-specific plasma IgG facilitates a focused examination of the most highly abundant plasma antibodies that are likely critical for endpoint neutralization of the parasite in the bloodstream and 2) understanding the contribution of plasma antibody binding to non-neutralizing epitopes of PfRH5 provides important knowledge for improved vaccine design. In this study, we establish that highly abundant non-neutralizing IgG lineages are prevalent within each donor’s PfRH5-specific plasma IgG repertoire. This was highlighted in Donor A, in which all top 6 IgG lineages (comprising ∼64% of circulating anti-PfRH5 IgG) are non-neutralizing individually. Several plasma mAbs bound linear epitopes such as the N-terminus, C-terminus, and IDL, each of which has been shown to be a target of non-neutralizing (GIA-negative) antibody responses to RH5.^29,35^ Further, it has been previously noted that anti-N-terminus IgG titers correlate with IgG serum titers against full length PfRH5.^25^ Consistent with this finding, N-terminus-directed antibodies were identified as highly abundant plasma IgG in an additional two donors in this study (pR26 and pR31, ranked 1^st^ and 6^th^ in abundance respectively). The immunodominance of these non-neutralizing linear epitopes within the vaccine-induced plasma IgG response to RH5.1 emphasizes the importance of ongoing immunogen design efforts^35^ in which these epitopes are either masked or removed. The PfRH5-specific plasma IgG response exhibited non-random germline gene use. IGHV5-51 was enriched among abundant, non-neutralizing antibodies targeting linear or disordered PfRH5 regions, consistent with antigen-driven biases toward frameworks permissive for flexible epitope recognition.^42^ In contrast, synergistic community 4 antibodies recurrently paired with IGLV3-21, as previously observed in RH5-specific mAbs,^29^ supporting a germline contribution to antibody synergy.

To dissect the functional implications of the polyclonal antibody response to RH5.1 vaccination, we employed Oligoclonal Reconstitution Experiments (OREs). These experiments allow us to create simplified pools of polyclonal antibodies, enabling detailed analysis of complex interactions within the plasma antibody response. This study marks the first use of OREs to investigate plasma antibody dynamics, providing a clearer understanding of how different antibody populations interact to influence overall GIA potency. Through OREs, we identified specific combinations of highly abundant antibodies in plasma that exhibit varied synergistic effects, which are critical to understanding the nuanced roles of various plasma-resident antibodies. The ORE analyses revealed that non-neutralizing plasma IgG targeting community 4 epitopes play a crucial role in enhancing the neutralization capacity of the polyclonal plasma antibody response. Individually, these antibodies do not inhibit parasite growth, yet their combined presence significantly improves overall neutralization potency. Synergistic neutralizing antibodies have been identified previously in B cell studies for a number of infectious diseases, including SARS-CoV,^43,44^ SARS-CoV-2,^45^ Coxsackievirus,^46^ HIV,^47^ HCV,^48^ and malaria.^36,49,50^ Further, non-neutralizing antibodies that synergize with neutralizing antibodies have also been demonstrated.^29,30,51^ However, here we show for the first time that, 1) synergistic antibodies can be detected circulating at high concentrations within the blood; 2) these synergistic antibody pairs can be detected circulating within a single donor; 3) synergistic antibody interactions can involve pairs of non-neutralizing antibodies; and 4) synergistic antibodies can be convergent, recurring features in response to some antigens.

The implications of these findings are substantial not only for PfRH5 vaccine development, but as a general paradigm shift in how we evaluate antibody responses to infection and vaccination. Here, by identifying key epitopes that facilitate synergistic interactions, we were able to confirm key contributions to GIA potency provided by non-neutralizing antibodies. The ORE approach serves as a powerful tool for unraveling the intricacies of polyclonal antibody responses within the circulating plasma, informing on key elements of vaccine design. In conclusion, this study offers a comprehensive analysis of the PfRH5 plasma antibody response and highlights the predominance of non-neutralizing antibodies that play a key role in polyclonal plasma neutralization. Through OREs, we have demonstrated the significance of synergistic interactions between antibodies targeting community 4a and 4d epitopes, emphasizing their critical role in the antibody response induced by the RH5.1 vaccine candidate. Importantly, these results are specific to UK adults, and similar repertoire analyses should be extended to more relevant populations such as African children. In total, these findings underscore the importance of considering antibody synergy in vaccine development, providing a foundation for engineering more effective immunogens against malaria and other infectious diseases.

## Supplementary information for

**Figure S1.**
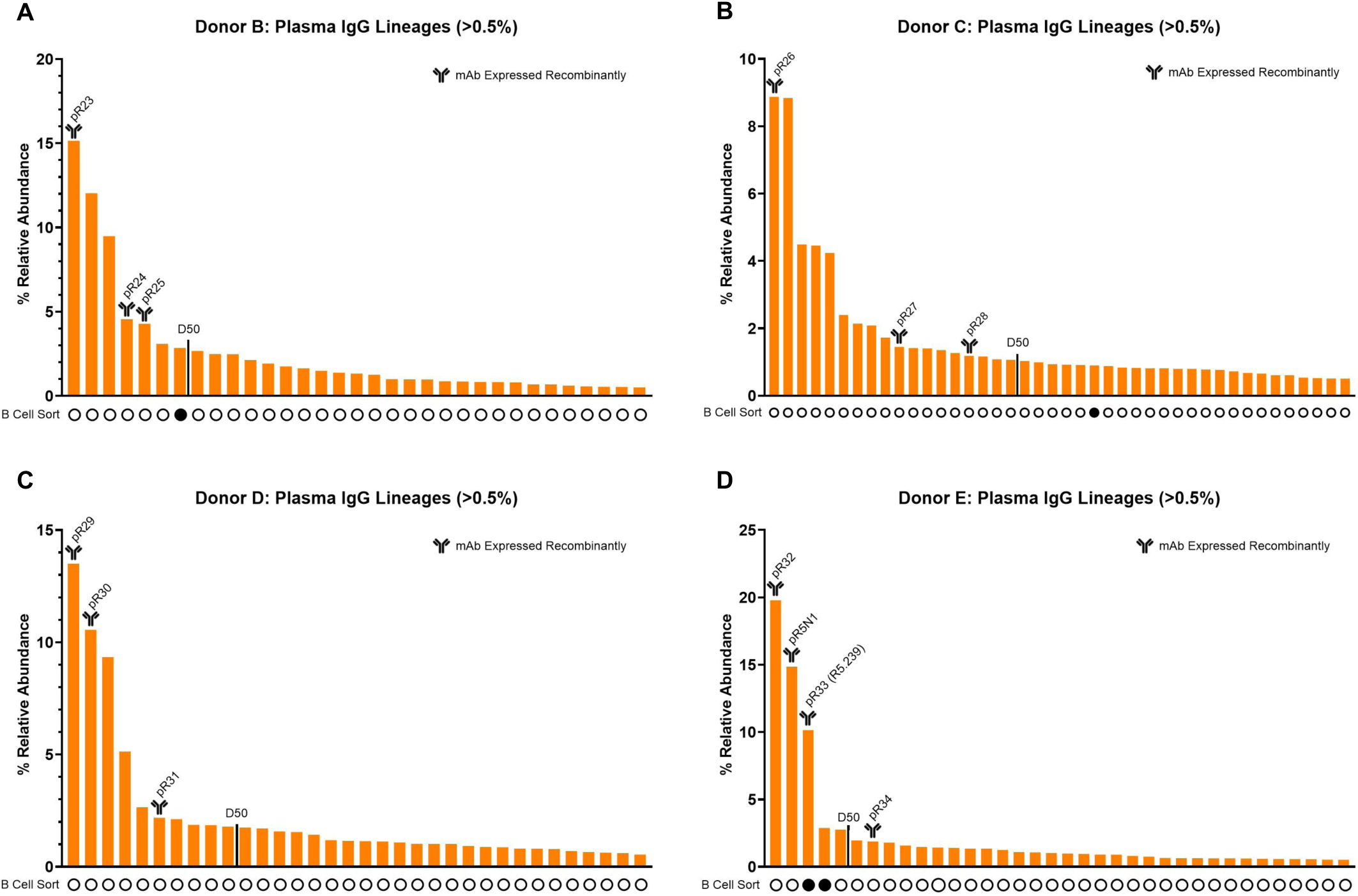
**The Anti-RH5.1/AS01_B_ Plasma IgG Repertoire of Four Donors**. Donors are labeled “B,” “C,” “D,” and “E” (Donor A is shown in Fig. 1B). Anti-RH5.1 IgG lineages ³0.5% relative abundance in (A-D) Donors B-E. The number of lineages that account for ∼50% of the anti-RH5.1 plasma repertoire is denoted by a line labeled with “D50.” Below each bar, a filled in circle acknowledges that the lineage CDRH3 has been identified prior using RH5-specific B-cell sorting.^29^

**Figure S2.**
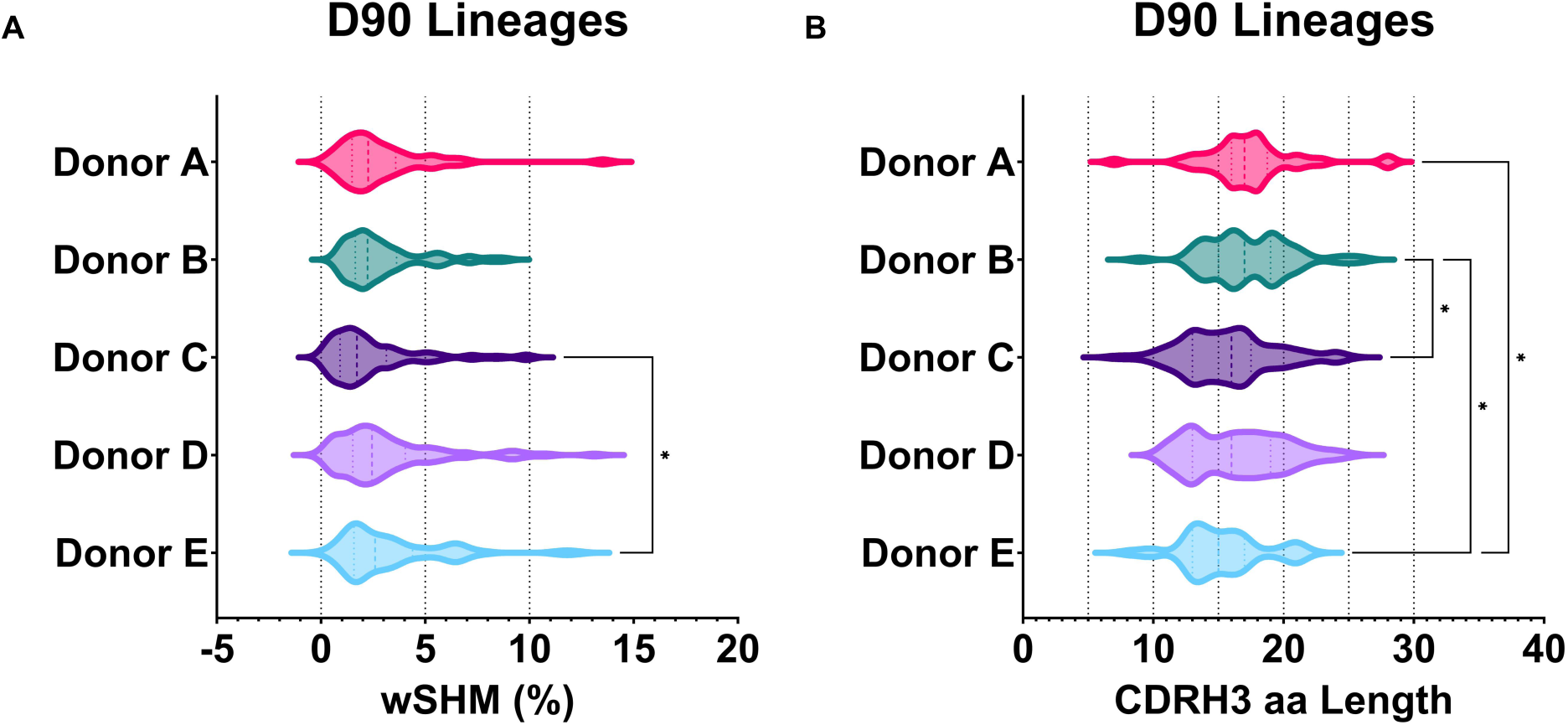
D90 BCR-Seq Features. (A) Violin plot of weighted SHM (wSHM) of top 90% relative abundance of lineages (D90) identified in all five donors. (B) CDRH3 length of D90 lineages identified in all five donors. (A-B) Dashed lines within violin plots represent different quartiles. * P < 0.05 ** P < 0.01, *** P < 0.001; **** P < 0.0001, determined by Kruskal-Wallis test with Dunn’s multiple comparison post-test.

**Figure S3.**
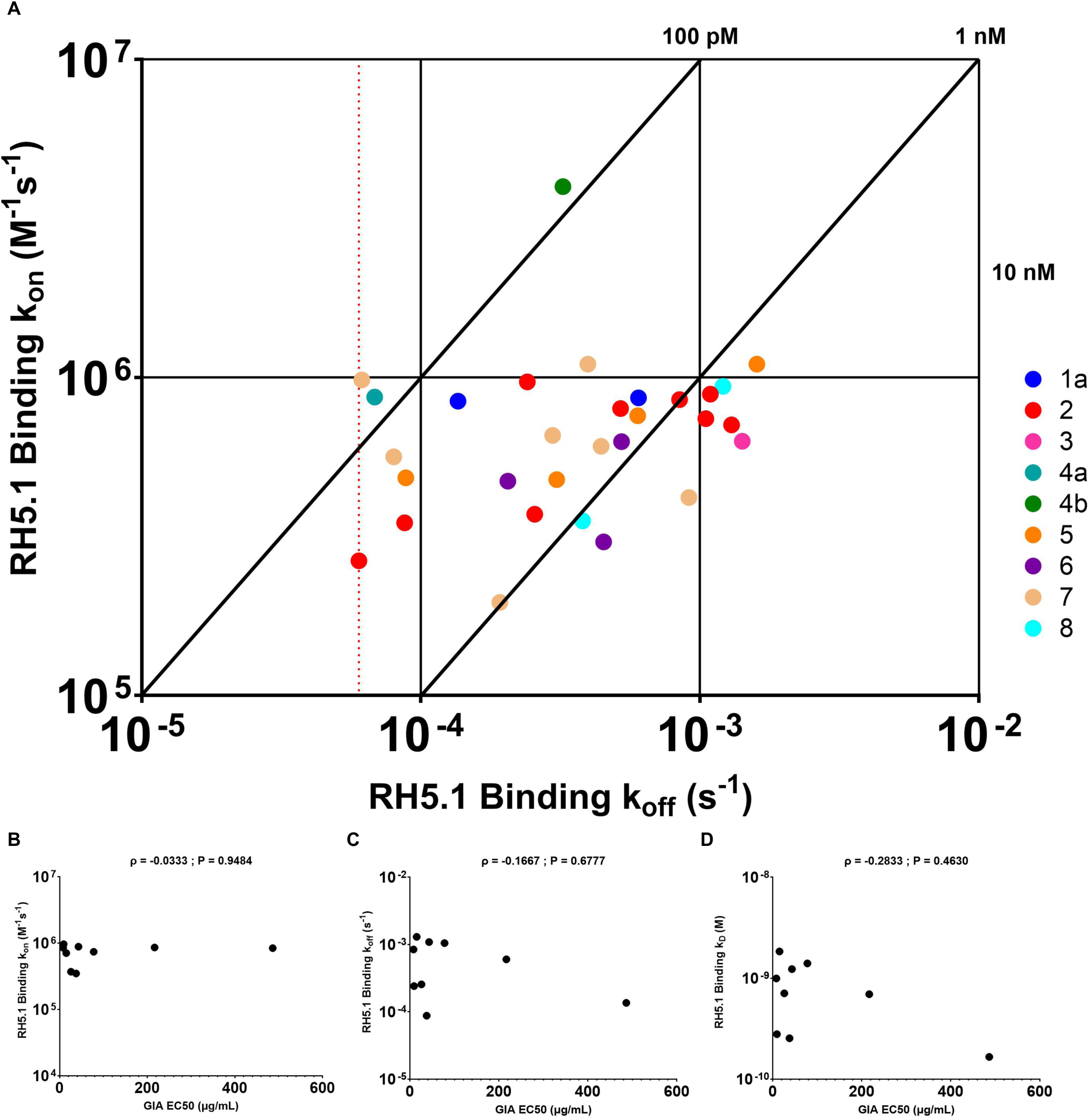
Binding Kinetics. (A) Iso-affinity plot showing kinetic rate constants for binding of mAbs to RH5.1 as determined by HT-SPR across all five donors. Diagonal lines represent equal affinity (K_D_ = k_off_/k_on_). Red dotted line indicates lowest limit of k_off_ measurement (6 x 10^-5^ s^-1^). mAbs are colored by HT-SPR epitope communities (see Fig. 2A and S4). Of the mAbs considered GIA-positive (≥20% GIA), correlations were determined for (B) k_on_ vs GIA EC_50_, (C) k_off_ vs GIA EC_50_, and (D) K_D_ vs GIA EC_50_. Spearman’s rank correlation coefficients (ρ) and two-tailed P values are shown.

**Figure S4.**
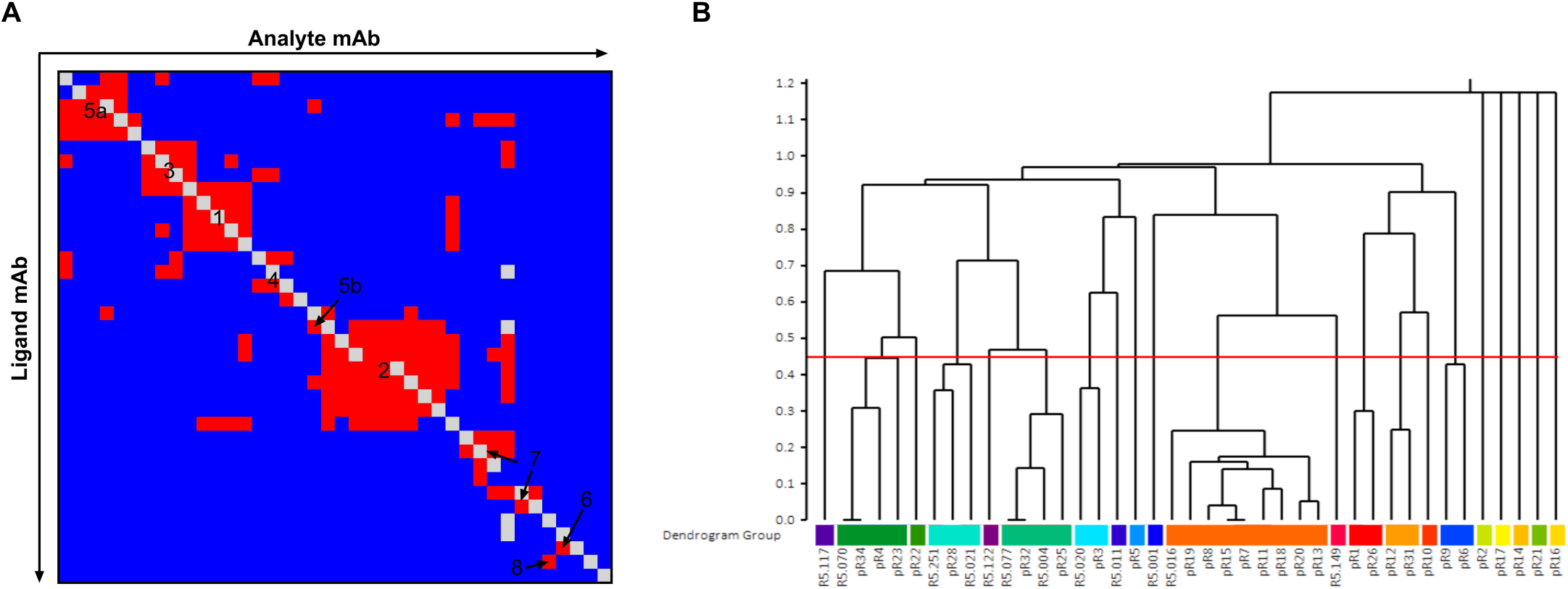
**LSA Carterra HT-SPR Analysis**. (A) Normalized RU values from competitive HT-SPR were automatically sorted into epitope communities using the Carterra Epitope software. Red squares indicates competition between mAbs, blue indicates no competition, and grey indicates values that were not measured or indicate self competion. (B) Dendrogram based on McQuitty clustering of competition data in (A). Communities were determined using a threshold height set by the behavior of known communities of control antibodies. ^29,30^

**Figure S5.**
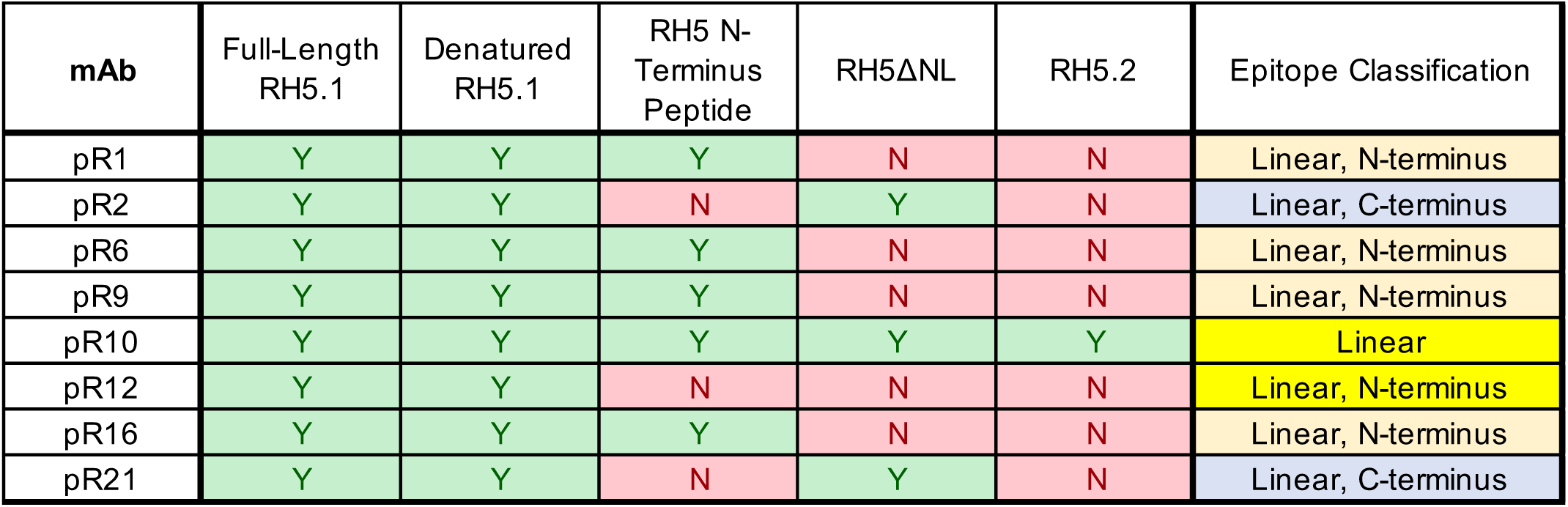
PfRH5 Variant ELISAs. mAbs belonging to new epitope communities 7, 8, or unknown communities were tested against PfRH5 variants, including full-length RH5.1, denatured RH5.1, PfRH5 N-terminus peptide, RH5ΔNL, and RH5.2. ELISA response to the RH5 N-terminus peptide confirmed community 7 as describing N-terminus-specific mAbs. pR12 bins with the remaining N-terminus mAbs despite not responding via ELISA. pR2 and pR21 have consistent binding patterns to RH5ΔNL, which contains the PfRH5 C-terminus epitope, and RH5.2, which does not. Given their specificity to denatured RH5.1 and RH5ΔNL and lack of binding to RH5.2, both were binned as C-terminus-specific mAbs.

**Figure S6.**
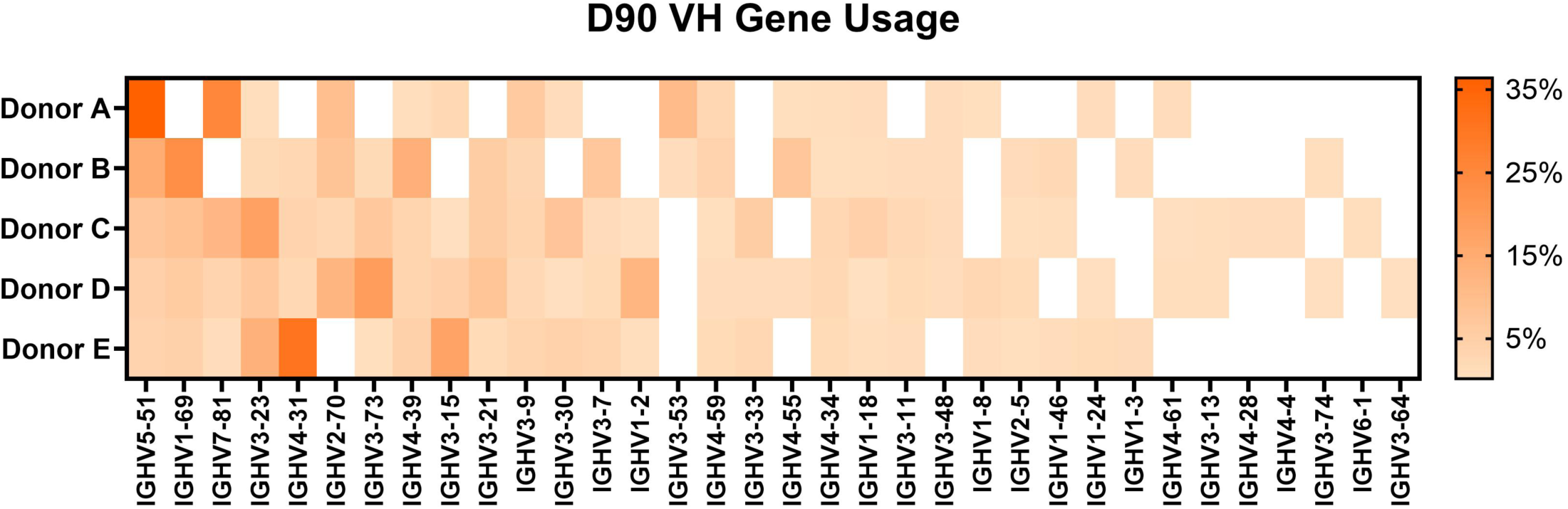
VH Gene Usage. VH gene usage data obtained from BCR-Seq in Donors A-E. For each VH gene, usage was summed as the percent relative abundance of individual lineages (as determined by Ig-Seq), which was normalized by the sum of all D90 lineage percent relative abundance values. The color bar on the right represents the normalized percent relative abundance of each VH gene per donor.

**Figure S7.**
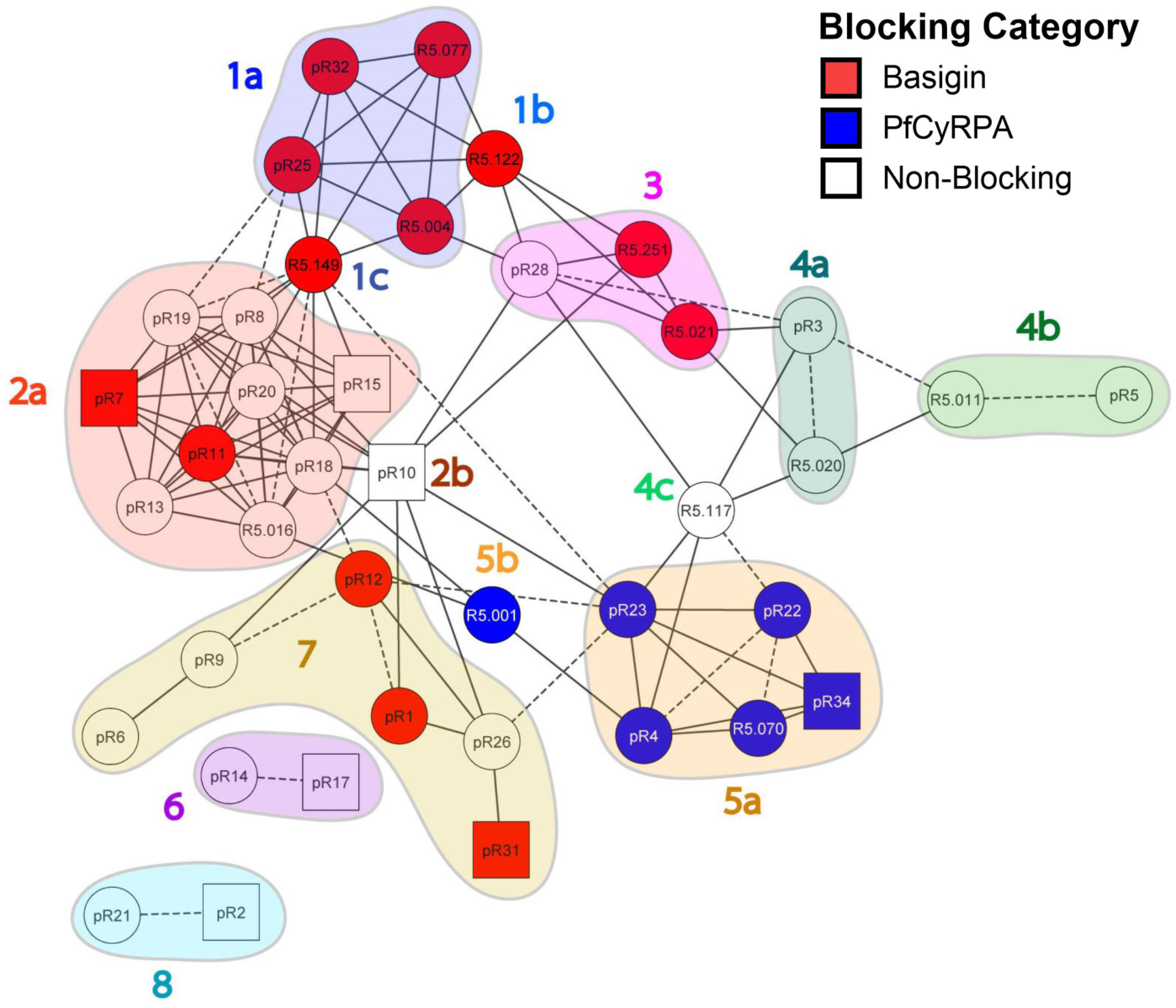
Basigin and PfCyRPA-Blocking Activity. Plasma mAb blocking activity of RH5.1 binding to (A) Basigin and (B) PfCyRPA, as determined by HT-SPR. Blocking threshold based upon positive controls R5.015 (PfCyRPA-blocking mAb), R5.004 (Basigin blocking mAb), and R5.011 (non-blocking mAb). Blocking data is superimposed onto community network plot in Fig. 2A.

**Figure S8.**
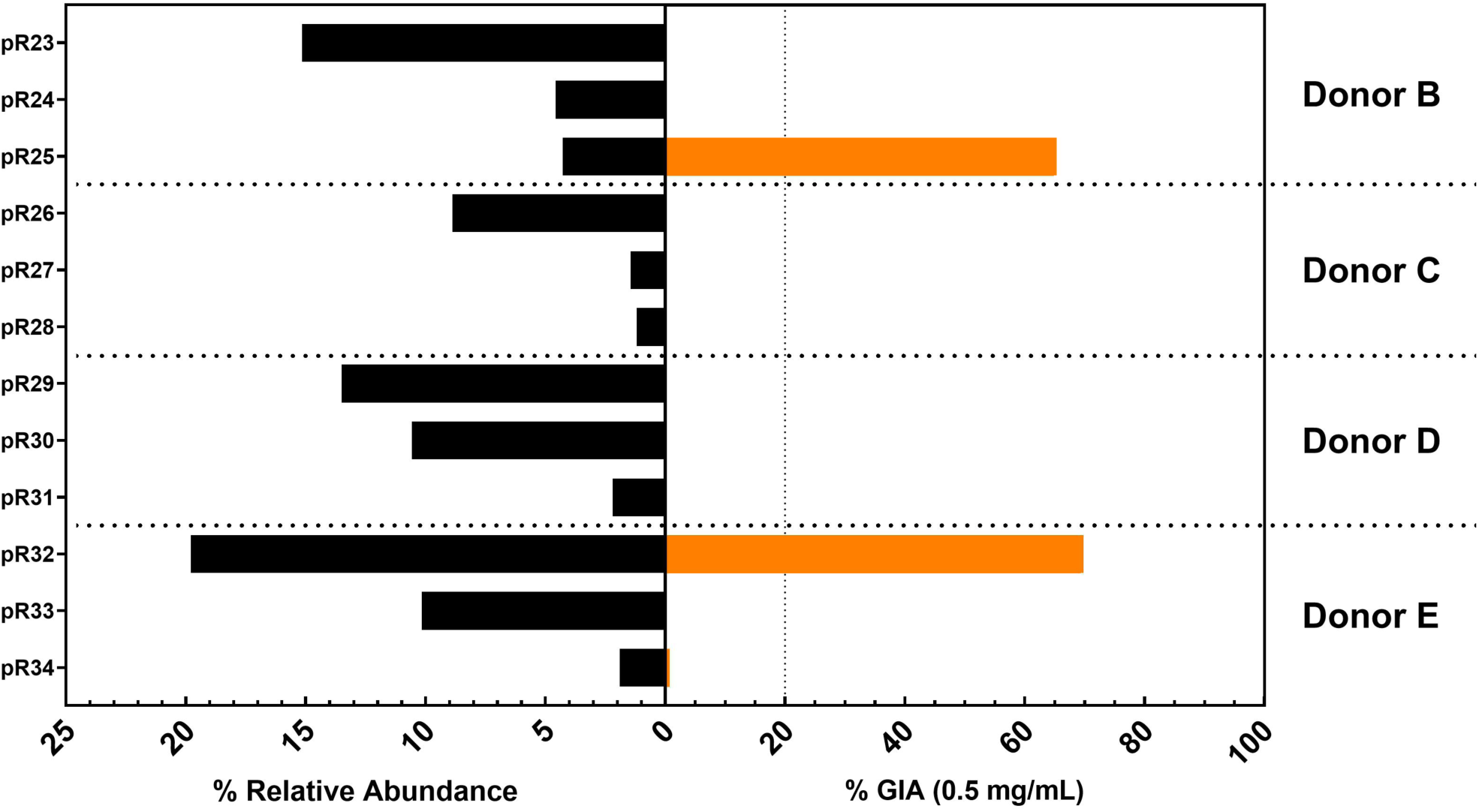
**Relative Abundance of Plasma mAbs versus GIA**. Data are shown for Donors B-E (Donor A is shown in Fig. 2D). pR23-25 for Donor B; pR26-28 for Donor C; pR29-31 for Donor D; and pR32-34 for Donor E. All mAbs were tested for GIA at 0.5 mg/mL.

**Figure S9.**
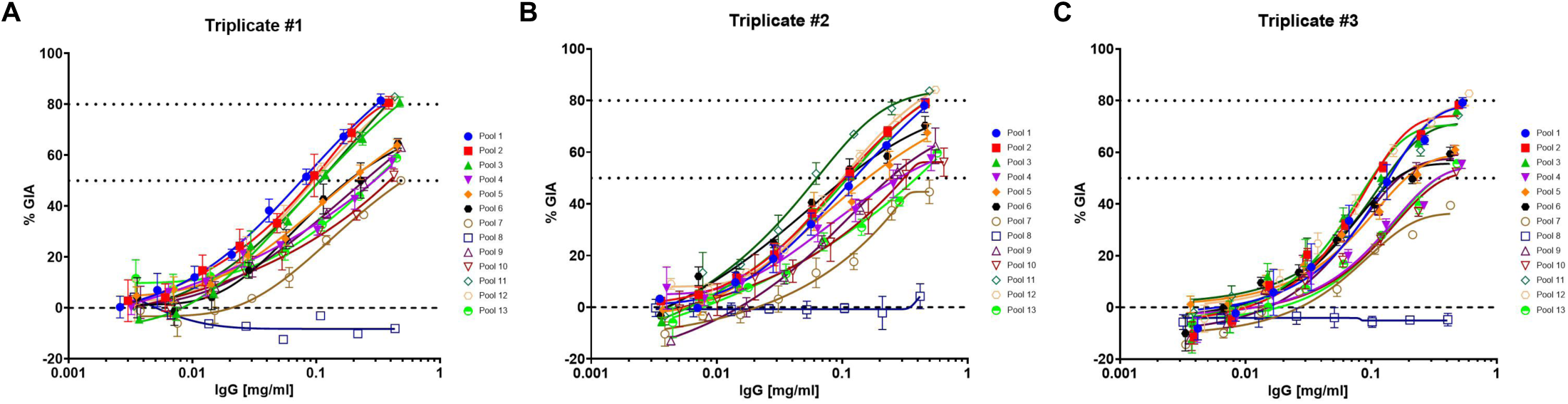
**ORE Pool Triplicates**. GIA dilution curves of Pools 1-13, starting at 0.5 mg/mL with an 8-step, 2-fold dilution series. Erythrocytes from distinct blood donor volunteers were used to validate results across three triplicate experiments. Pools #1-13 for (A) triplicate experiment #1, (B) triplicate experiment #2, and (C) triplicate experiment #3.

**Figure S10.**
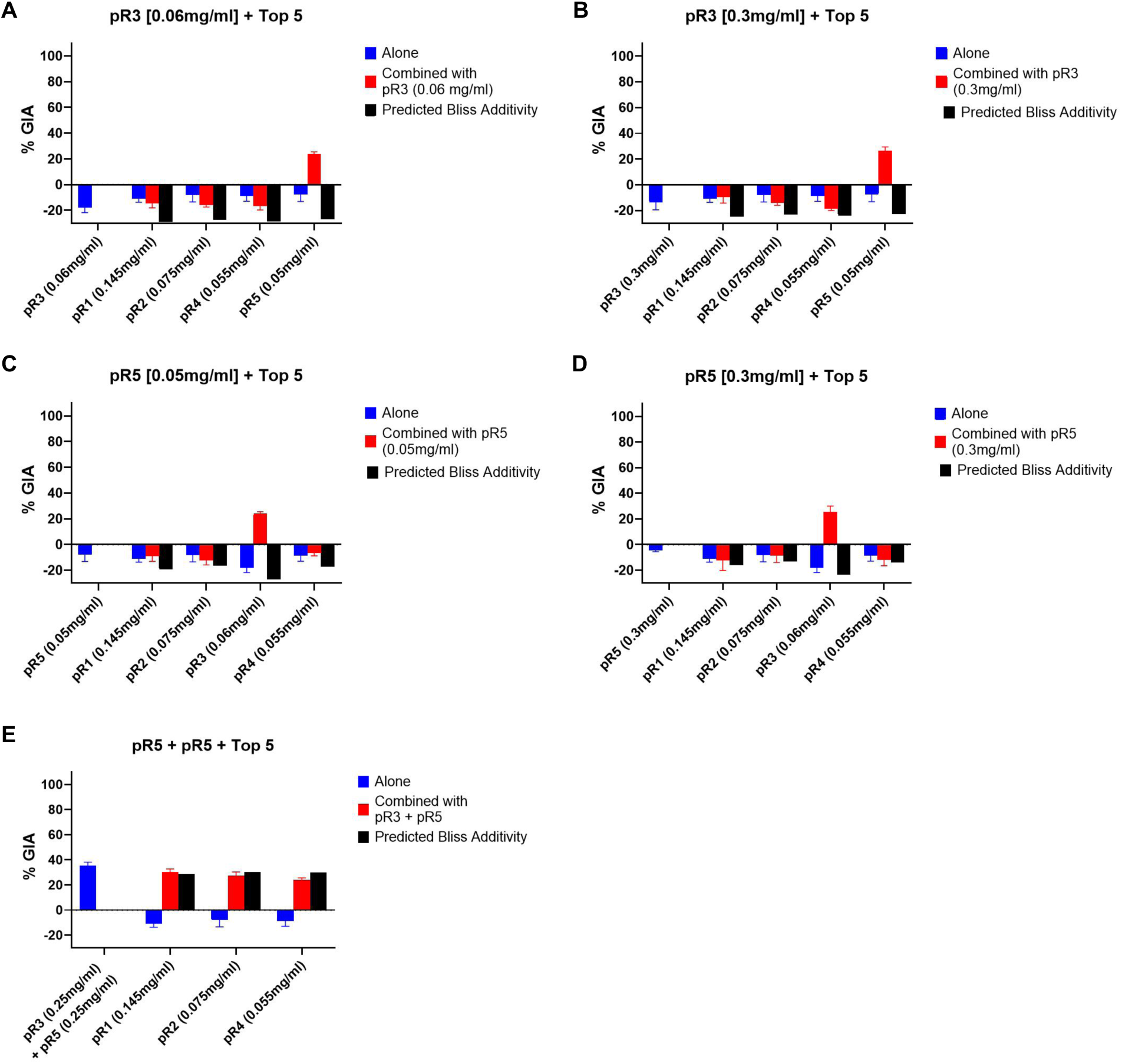
**Synergy Screens Among Top 5 Lineage mAbs**. pR1, pR2, and pR4 were held at concentrations representing their normalized relative percent abundance within a 0.5 mg/mL pool. They were mixed with either pR3 and/or pR5. pR3 was tested at (A) its relative pool concentration (0.06 mg/mL), or (B) in excess at 0.3 mg/mL. pR5 was tested at (C) its relative pool concentration (0.05 mg/mL) or (D) in excess at 0.3 mg/mL. (E) pR5.003 and pR5.005 together were tested in excess at 0.25 mg/mL each.

**Figure S11.**
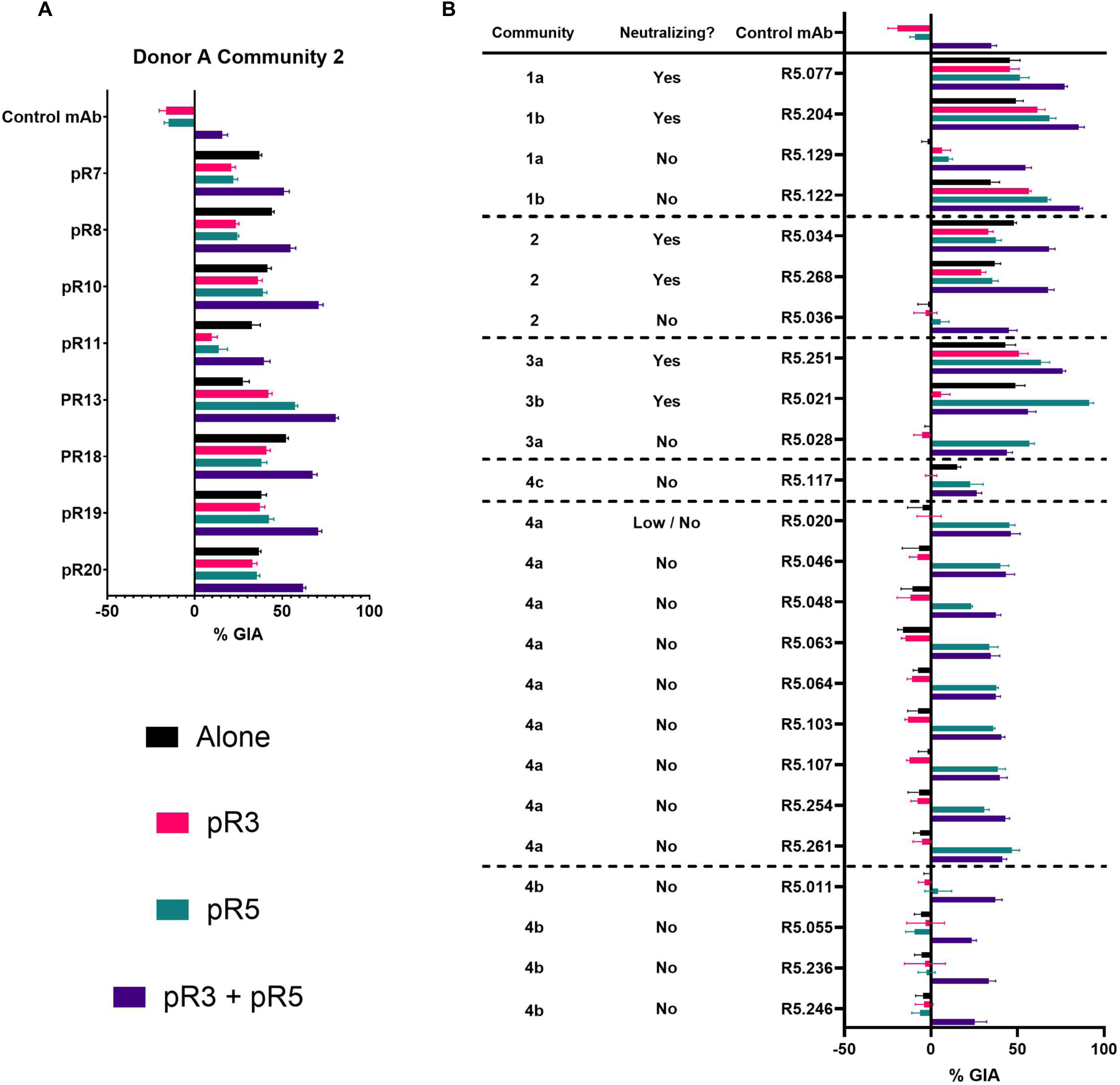
(A) Synergy GIA screens of community 2 mAbs used in OREs in Figure 2 with either Community 4a pR3 (0.2 mg/mL), Community 4b pR5 (0.2 mg/mL), or both (0.4 mg/mL combined). All test mAbs were held at their EC_50_ concentration except pR10 and pR13, which were held at 0.5 mg/mL. (B) Synergy GIA screens of previously defined neutralizing and non-neutralizing mAbs^29,30^ with either Community 4a pR3, Community 4b pR5, or both. Dashed lines indicate grouping of separate epitope communities. Neutralizing mAbs were tested at their respective EC_50_ values, while non-neutralizing mAbs were tested at 0.5 mg/mL.

**Figure S12.**
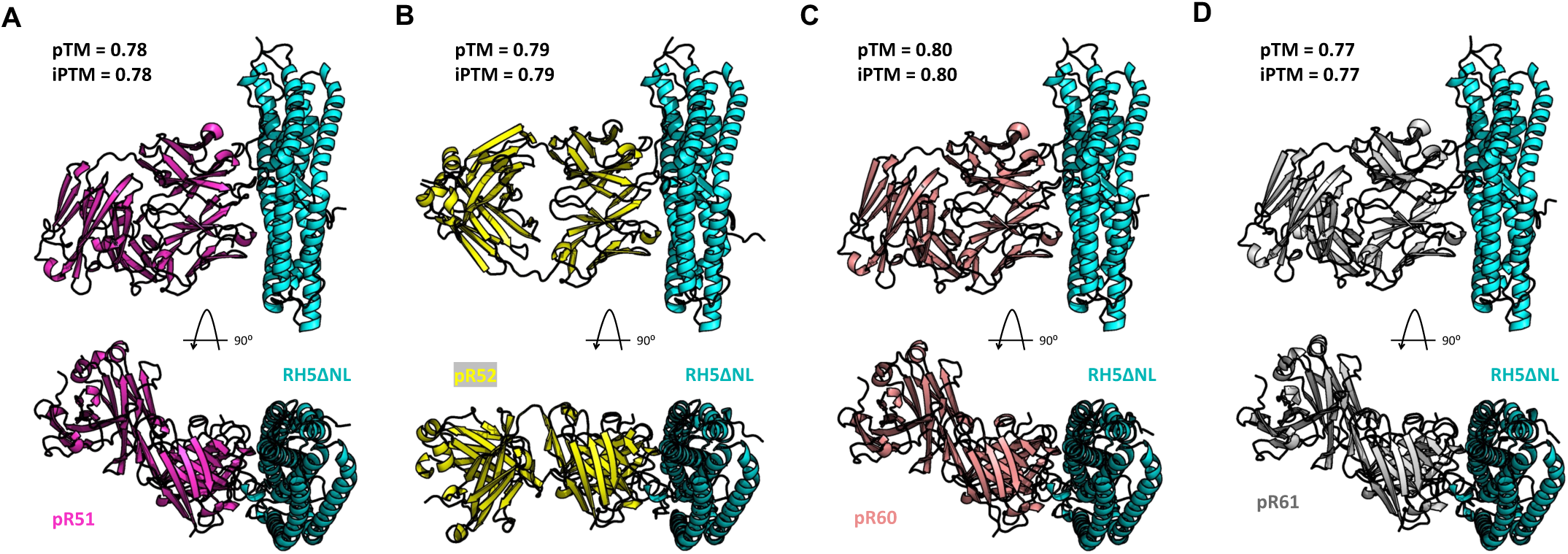
**AlphaFold 3 Predictions**. Front and top view of AlphaFold 3 predictions of the interface between RH5ΔNL and Fab fragments of mAbs A) pR51, B) pR52, C) pR60, and (D) pR61. pTM and iPTM scores are listed for each fold. All other Fabs resulted in poor fold scores.

**Figure S13.**
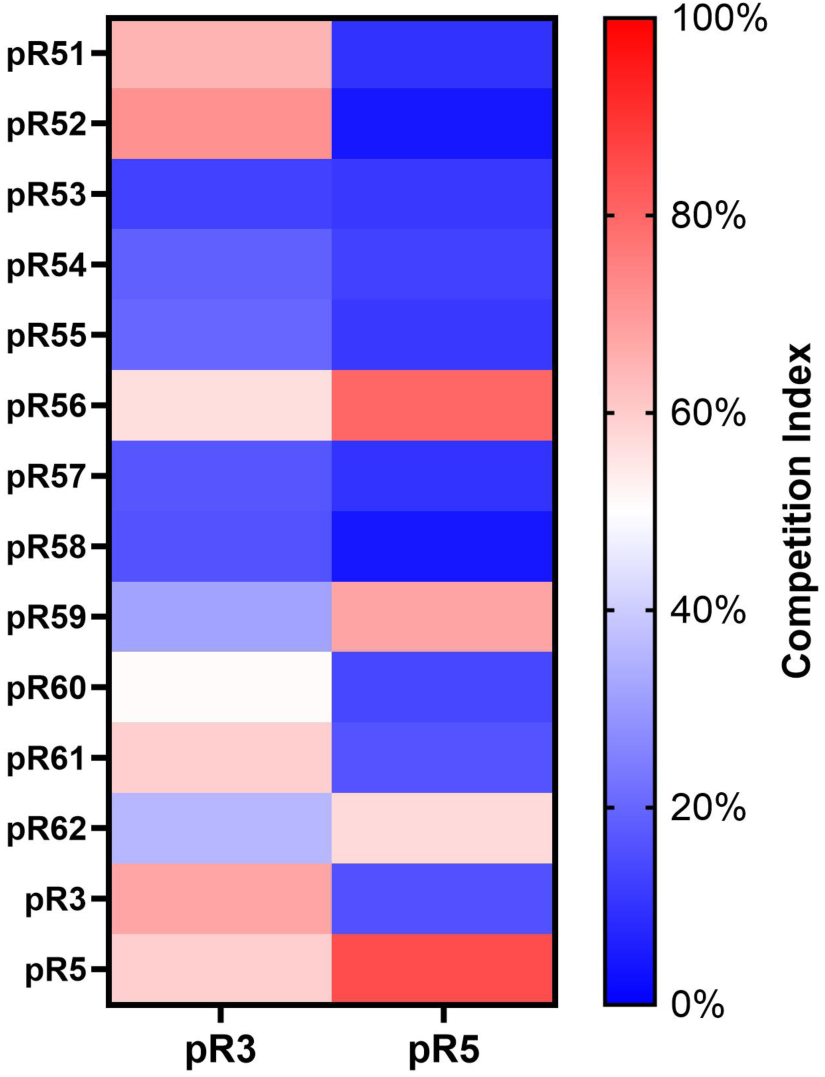
**Indirect Competition ELISA with pR3 and pR5**. Test mAbs pR51-pR62, alongside pR3 and pR5 were tested for competition against RH5.1 with pR3 or pR5. The competition index is calculated by normalizing all OD values to the highest OD value recorded (no competing mAb control), inverting these values, and converting them to percentages. This is visualized in a color gradient, with from blue (0%) to white (50%) to red (100%). A competition index ≥50% indicates the mAb competed with pR3 or pR5.

**Figure S14.**
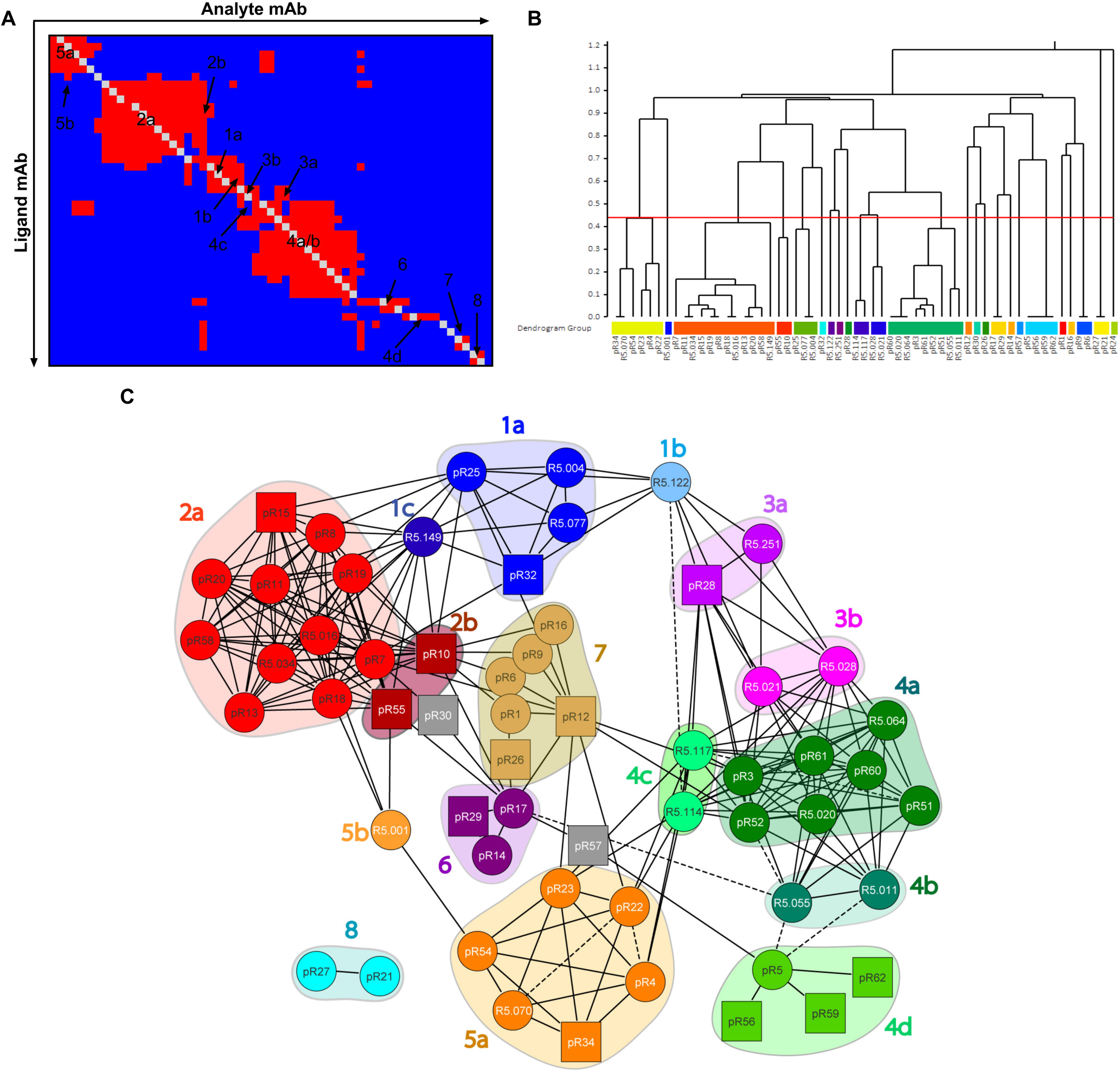
**HT-SPR of Putative Community 4 mAbs**. (A) Normalized RU values from competitive HT-SPR were automatically sorted into epitope communities using the Carterra Epitope software. Red squares indicates competition between mAbs, blue indicates no competition, and grey indicates values that were not measured or indicate self-competition. (B) Dendrogram based on McQuitty clustering of competition data in (A) Communities were determined using a threshold height set by the behavior of known communities of control antibodies.^29,30^ (C) Pairwise competition among plasma mAbs, designated from (A-B). Each community is identified by a color and number code, and subcommunities indicated by a letter. Individual mAbs are represented by nodes. Control mAbs^29,30^ are listed with the prefix “R5,” while plasma mAbs are listed with the prefix “pR.” Dashed lines and solid lines indicate unidirectional and bidirectional competition, respectively. Square nodes indicate that a mAb was excluded as either ligand or analyte. mAbs pR24, pR30, and pR57 do not fall into pre-defined communities and are not assigned a label.

## Methods

### Donor cohort & collection of blood plasma and PBMCs

Donor plasma and peripheral blood mononuclear cells (PBMCs) were isolated from adult human volunteers immunized with the RH5.1/AS01_B_ vaccine candidate during the VAC063 clinical trial.^27^ VAC063 was an open-label, non-randomized, multi-center, dose-escalation Phase 1/2a trial designed to assess the safety, immunogenicity, and efficacy of the *P. falciparum* blood-stage malaria vaccine candidate RH5.1/AS01_B_. Volunteers were healthy, malaria-naïve UK adults aged 18-45 years. The volunteers evaluated in the study reported here belonged to Group 5 (as reported in the original trial publication^27^) and received three doses of 10 µg RH5.1 in 0.5 mL AS01_B_ (GlaxoSmithKline), each dose spaced 28 days apart, followed by an intravenous blood-stage CHMI at day 70 (14 days post-3rd immunization).

The VAC063 study was conducted in the UK at the Centre for Clinical Vaccinology and Tropical Medicine, University of Oxford, Oxford; Guys and St Thomas’ NIHR CRF, London; and the NIHR Wellcome Trust Clinical Research Facility in Southampton. The study received ethical approval from the UK NHS Research Ethics Service (Oxfordshire Research Ethics Committee A, Ref 16/SC/0345) and was approved by the UK Medicines and Healthcare products Regulatory Agency (Ref 21584/0362/001-0011). Volunteers signed written consent forms and consent was verified before each vaccination. The trial was registered on ClinicalTrials.gov (NCT02927145) and was conducted according to the principles of the current revision of the Declaration of Helsinki 2008 and in full conformity with the ICH guidelines for Good Clinical Practice.

Donations of human red blood cells (RBCs) from healthy adult volunteers for use in assays received ethical approval from the UK NHS Research Ethics Service (London – City & East Research Ethics Committee, Ref 18/LO/0415).

Human blood samples were collected into lithium heparin-treated vacutainer blood collection systems (Becton Dickinson). PBMCs were isolated 7 days following 3rd immunization (day 63) and frozen in fetal calf serum containing 10% dimethyl sulfoxide and stored in liquid nitrogen.

Plasma samples were isolated 13 days following 3rd immunization / 1 day prior to CHMI (day 69) and stored at −80°C.

### Expression and purification of protein

RH5.1: RH5.1 used in Ig-Seq and additional assays was provided by The Draper Lab (University of Oxford). RH5.1 is a variant of full-length 3D7 clone *P. falciparum* RH5 antigen (amino acids E26-Q526) that contains Thr to Ala mutations at all four putative N-linked glycosylation sequons (N-X-S/T).^52^ As described previously, RH5.1 was expressed in *Drosophila* S2 cells and purified using C-tag technology to meet current Good Manufacturing Practice (cGMP) compliance.^26^ RH5ΔNL: RH5ΔNL is a variant of full-length 3D7 clone *P. falciparum* Rh5 antigen that excludes the N-terminus and IDL epitopes, specifically residues K140-K247 and N297-Q526 with two mutations to delete two N-linked glycosylation sites (T216A and T299A). As described previously, RH5ΔNL was expressed in *Drosophila* S2 cells.^30^

RH5 N-terminus: The peptide encodes 3D7 clone residues F1-K116 and was chemically synthesized, as described previously.^53^

RH5.2: RH5.2 contains 18 stabilizing mutations: I157L, D183E, A233K, M304F, K312N, L314F, K316N, M330N, S370A, S381N, T384K, L392K, T395N, N398E, R458K, N463K, S467A, F505L. It was expressed in *Drosophila* S2 lines, as described previously.^35,54^

Basigin: Recombinant basigin was expressed through transient transfection of Expi293 cells and contained native residues M1-L206, followed by rat CD4 domains 3 and 4 and a C-terminal hexahistidine tag. Basigin was purified from the supernatant by immobilized metal affinity chromatography using a Ni^2+^ resin followed by size-exclusion chromatography (SEC) into TBS pH 7.4 (20 mM TRIS-HCl, 150 mM NaCl) as previously described.^14^

PfCyRPA: Recombinant PfCYRA was used in competitive surface plasmon resonance (SPR). Full-length PfCyRPA (residues 29-362) was expressed in Expi293 cells and purified through C-tag affinity chromatography follow by SEC into TBS pH 7.4 as previously described.^36^

### VH repertoire sequencing (BCR-Seq)

PBMCs were pelleted at 300 x g for 10 min, followed by resuspension in TRIzol Reagent (Thermo Fisher Scientific; Cat: 15596026) to lyse cells and release RNA. Total RNA was extracted using the RNeasy Mini Kit (Qiagen; Cat: 74104) according to the manufacturer’s protocol. First-strand cDNA was synthesized from 500 ng of isolated mRNA using SuperScript IV (Thermo Fisher Scientific; Cat: 18090010). cDNA encoding the VH regions of IgG, IgA, and IgM repertoires was amplified with a multiplex primer set^55^ using the FastStart High Fidelity PCR System (Sigma-Aldrich; Cat: 3553400001) under the following conditions: 95 °C for 2 min; [92 °C for 30 s, 50 °C for 30 s, 72 °C for 1 min] x 4 cycles; [92 °C for 30 s, 55 °C for 30 s, 72 °C for 1 min] x 4 cycles; [92 °C for 30 s, 63 °C for 30 s, 72 °C for 1 min] x 22 cycles; 72 °C for 7 min; hold at 4 °C. The amplified product was PCR purified using DNA Clean & Concentrator (Zymo; Cat: D4030), fol-lowed by gel electrophoresis using a 1% agarose gel. Amplicons were extracted from gel and purified using Zymoclean Gel DNA Recovery Kit (Zymo; Cat: D4002). Products were sequenced by 2 x 300 paired-end Illumina MiSeq by the Genome Sequencing and Analysis Facility (GSAF) at The University of Texas at Austin (UT Austin).

### Paired VH:VL repertoire sequencing (BCR-Seq)

The paired VH:VL amplicon library was prepared using a flow-focusing device (FFD) which captures individual B cells in emulsion droplets, followed by an emulsion-based overlap extension reverse-transcriptase-PCR (OE RT-PCR) designed to link native VH and VL cDNA, from which amplicons were prepared and sequenced from subsequent PCR reactions, as previously described.^56^ To summarize, 2 x 10^6^ PBMCs were co-emulsified with magnetic Dynabeads Oligo(dT)_25_ (Thermo Fisher Scientific; Cat: 61002) in lysis buffer (100 mM Tris pH 7.5, 500 mM LiCl, 10 mM EDTA, 1% lithium dodecyl sulfate, and 5 mM dithiothreitol) using the FFD. Captured magnetic bead emulsions were broken using hydrated ether, followed by magnetic bead washing and resuspension in a one-step OR RT-PCR solution containing overlap extension VH and VL primer sets, as previously described.^56^ Magnetic beads resuspended in this mixture were re-emulsified in a dispersion tube (IKA; Cat: 000370310) and subjected to OE-RT PCR under the following conditions: 68 °C for 30 min; 94 °C for 2 min; [94 °C for 30 s, 60 °C for 30 s, and 68 °C for 2 min] x 25 cycles; 68 °C for 7min; hold at 4 °C. Following the PCR, emulsions were broken using hydrated ether and amplicons were extracted and purified using DNA Clean & Concentrator (Zymo; Cat: D4030). Product was further amplified and barcoded using nested PCRs and MiSeq PCRs, as previously described.^57^ Products were sequenced by 2 x 300 paired-end Illumina MiSeq by the GSAF at UT Austin.

### Bioinformatics analyses

Bioinformatics analysis of BCR-Seq (VH repertoire sequencing, paired VH:VL repertoire sequencing) was completed, as described previously.^58^ From VH repertoire sequencing, paired-end FASTQ files were quality filtered and merged using PEAR software,^59^ followed by IMGT VDJ germline alignment using MiXCR software.^60^ VH sequences were clustered into sets of unique lineages using a single-linkage hierarchical clustering algorithm based on 90% identify measured by Levenshtein distance across the CDRH3 amino acid sequences. From paired VH:VL repertoire sequencing, paired-end FASTQ files were quality filtered using Trimmomatic^61^ software, followed by IMGT VDJ germline alignment using MiXCR software.^60^ Productive VH:VL paired reads were clustered by 90% CDRH3 homology using UCLUST^62^ software.

### Bottom-Up LC-MS/MS Sample Processing (Ig-Seq)

To purify total IgG, 1 mL of plasma was diluted 1:1 with Dulbecco’s PBS 1X and applied to a Protein G Plus Agarose (Thermo Scientific; Cat: 22852) affinity chromatography column. IgG was eluted using 100 mM glycine-HCl, pH 2.7 and immediately neutralized using 1 M Tris-HCl, pH 8.0. Neutralized IgG was concentrated and buffer-exchanged into Dulbecco’s PBS 1X using a Vivaspin® Turbo 4, 30 kDa (Sartorius; Cat: VS04T21). The total IgG was further cleaved into F(ab’)_2_ using IdeS at 37 °C for 2 h.

RH5.1-specific F(ab’) _2_ was isolated sequentially by affinity chromatography as follows. First, 1 mg recombinant RH5.1 antigen and 0.05 mg dry NHS-activated agarose resin (Thermo Scientific; Cat: 26196) were incubated with spinning at 4 °C O/N, followed by blocking using 1M ethanolamine, pH 8.2 with spinning at r.t for 20 min. The antigen-conjugated resin was loaded into 0.5 mL spin columns, washed 12 times with 0.4 mL Dulbecco’s PBS by centrifuging 1000 x g for 30 s, and mixed with F(ab’)_2_ (variable quantity, depending on donor titer) with spinning at room temperature (r.t.) for 1 h. Following incubation, the antigen-F(ab’)_2_-resin was loaded into 0.5 mL spin columns again and the flow-through was captured by centrifuging 1000 x g for 30 s. At least 10 μg of flow-through was saved for bottom-up proteomics processing. The column was washed 12 times again with 0.4 mL Dulbecco’s PBS via centrifugation and F(ab’)_2_ was eluted by centrifuging 1000 x g for 30 s in 0.36 mL fractions using 1% formic acid (FA). Immediately after each elution, 20 μL of the sample was neutralized with 30 μL of 1 M Tris-HCl, pH 8.5 to be tested via indirect ELISA and determine if antigen-specific F(ab’)_2_ had been successfully captured. Elutions determined to contain antigen-specific F(ab’)_2_ were speed-vacuumed at 45 °C for ∼1.5 h in aqueous mode until ∼5-10 µL of final volume remained. Concentrated eluates were combined into a single tube, 0.2 v/v 1 M Tris, pH 8.5 was added to the sample, and final pH was adjusted to 7-7.5 using 1 M Tris / 3 M NaOH at a final volume of 50 μL.

10 μg flow-through was adjusted to 50 μL final volume and processed with the elution in the following steps. The samples were denatured using an equal volume (50 µL) of 100% 2,2,2-trifluoroethanol (TFE) (Sigma-Aldrich; Cat: T63002-25G). Disulfide-bonds were reduced with 1% v/v (1 µL) of 500 mM Bond-Breaker Tris (2-carboxyethyl) phosphine hydrochloride (TCEP) solution (Thermo Fisher Scientific, MA; Cat: 77720) and incubated at 55 °C for 1 h in a dry shaker. The reduced samples were alkylated by incubating with ∼3% v/v (3 µL) of 550 mM iodoacetamide (IAM) (Sigma-Aldrich Cat: I1149-5G) at r.t. for 30 min. Samples were diluted to 5% TFE, 0.5 mM TCEP, and 1.65 mM IAM by adding 40 mM Tris-HCl, pH 8.0 (892 µL). 2 µg MS grade Trypsin gold (Promega; Cat: V5280) was added to each sample and incubated at 37 °C for 4 h. The digestion reaction was quenched by adding 10 µL 100% FA.

Samples were speed-vacuumed at 45 °C for ∼3 h until less than 20 µL was left. Dried samples were resuspended in 100 µL Buffer A (5% HPLC-Grade ACN, 95% HPCL-Grade water, 0.1% FA). Samples were desalted using the following steps, all via centrifugation at 1000 x g for 30 s: HyperSep SpinTip SPE extraction C18 tips (1 per sample) (Thermo Scientific; Cat: 60109-412) were washed 3 times with 67 µL Buffer B (0.1% FA in HPLC-Grade water diluted with HPLC-Grade CAN at a 2:3 ratio) followed by 3 washes with 67 µL 0.1% FA. Samples were loaded onto C18 tips and centrifuged. C18 tips were washed 3 times with 67 µL 0.1% FA. C18 elutions were captured in triplicate 67 µL fractions using Buffer B. C18 elutions were combined and speed-vacuumed at 45 °C for ∼30 min until ∼5-10 µL of volume remained. C18 elutions were resuspended to 50 µL in Buffer A and submitted to the UT Austin Center for Biomedical Research Support Biological Mass Spectrometry Facility (RRID: SCR_021728) for protein identification by LC-MS/MS using a Thermo Ultimate 3000 RSLCnano ultra-high-performance liquid chromatography (UHPLC) coupled to a Thermo Scientific Orbitrap Fusion Tribrid mass spectrometer.

### Sample run method for LC-MS/MS

Peptide samples were dried, resuspended in 0.1% FA, and transferred to the LC instrument auto-sampler. Peptides were first loaded onto an Acclaim™ PepMap™ 100 C18 HPLC trap column (Thermo Scientific; Cat: 164535) for desalting and concentration, and subsequently separated on an EASY-Spray™ HPLC analytical column (Thermo Scientific; Cat: ES902). A gradient from 5 to 45% mobile phase B (0.1% FA in ACN) eluted peptides by increasing hydrophobicity over 100 min with a total run time of 120 min. MS data profile in positive mode was acquired in the Orbitrap with the following settings: 3 s cycle time, 120,000 resolution, scan range 400-1600 m/z, maximum injection time of 60 ms, and 60% RF lens value. A minimum signal intensity of 5 x 10^3^ was required to trigger a data-dependent scan with charge states 2 to 6 inclusive. The dynamic exclusion properties were set to allow each precursor to be selected up to two times (n=2) before being added to the dynamic exclusion list, with additional properties of ±25 ppm and 30 s exclusion. A targeted mass exclusion list containing the m/z values for 88 IgG peptides was included. Centroid MS/MS data were acquired in the linear ion trap using quadrupole isolation followed by higher-energy collisional dissociation (HCD) fragmentation with the following settings: 1.6 m/z isolation window, stepped HCD collision energies (27, 31, and 35%), and rapid ion trap scan rate with dynamic maximum injection time mode.

### MS search workflow and data analysis

Proteomics analyses following LC-MS/MS was performed as previously described.^31,55^ Per donor, a personalized MS search database was created using (i) the full-length RH5.1 sequence, (ii) individual VH and VL reads sequenced gathered from VH-only and paired VH:VL sequenced (BCR-Seq), and (iii) a common contaminants list provided by MaxQuant. Thermo Proteome Discover software 1.4 identified matching experimental spectra against each subject’s custom MS search database.

Following the search, MS data were analyzed as previously described. In brief, peptide abundance was measured using the sum of XIC precursor areas. We summed the XIC areas of anti-RH5.1 CDRH3 peptides that are (a) uniquely associated with a single lineage (as determined by BCR-Seq clustering), (b) that have peptide spectrum matches (PSMs) ≥ 2 across triplicate elution injections, and (c) are found >5-fold in the elution compared to the flow-through. Any CDRH3 peptides found in multiple clonotypes or having only 1 PSM detected across triplicate elution injections were excluded from subsequent clonotyping analyses. The relative abundance (%) of individual anti-RH5.1 lineages was calculated as the XIC area of a single lineage divided by the XIC area of all anti-RH5.1 lineages identified.

To compare the molecular features of the plasma antibody response, somatic hypermutation (SHM) of individual lineages was determined as a weighted average of the SHM for all VH reads clustered within that lineage, with weights based on the read count for each VH read. When determining CDRH3 length, CDRH3 annotation included “C” at the start of the sequence. When comparing VH usage, MiXCR^60^ software was used to determine VH gene assignment.

### Monoclonal antibody cloning

Cognate VH and VL antibody sequences of interest were synthesized and cloned into customized AbVec expression plasmids containing a human IgG1 Fc region (AbVec-HCIgG1, AbVec-LLC, AbVec-KLC, a gift from Patrick C. Wilson, University of Chicago, USA) by Twist Bioscience. Plasmid provided by Twist Bioscience was used in subsequent transfections.

### Expression and purification of IgG

Transfections of HEK293F cells using a 1:1 ratio of heavy chain (HC) and light chain (LC) plasmids were set up in 10 mL culture volumes in 50 mL vented cap reaction tubes using ExpifectamineTM transfection kit (Life Technologies) as per the manufacturer’s instructions. IgG purification was carried out using Econo-PAC® chromatography columns (Bio-Rad) and Protein A Sepharose (Sigma-Aldrich), and purified mAbs were buffer-exchanged into phosphate-buffered saline (PBS).

### Indirect ELISA

Nunc Maxisorp plates were coated overnight (>16 h) with RH5.1, RH5ΔNL, or RH5.2 at 2 μg/mL. For the ELISA against denatured antigen, RH5.1 protein was heated for 10 min at 80 °C using a PCR machine and coated on plates at 2 μg/mL. Peptide ELISAs were performed as previously described^29^ using streptavidin-coated plates. Plates were washed in wash buffer (PBS with 0.05 % Tween 20 [PBST]) and blocked with 100 μL/well of Blocker Casein (Thermo Fisher Scientific; Cat: 37528) for 1 h. Plates were washed and antibodies at 10 μg/mL diluted in casein were added.

Following a 2 h incubation, plates were washed and a 1:1000 dilution of goat anti-human IgG (γ-chain specific) alkaline phosphate conjugate antibody (Sigma-Aldrich; Cat: A3187) was added and incubated for 1 h. Plates were washed in washing buffer and 100 μL development buffer (p-nitrophenyl phosphate substrate diluted in diethanolamine buffer) was added per well and developed according to internal controls. All mAbs were tested in duplicate. Unless otherwise stated, 50 μL was added per well and all steps were carried out at r.t.

### Indirect competition ELISA

96-well EIA/RIA Clear Flat Bottom Polystyrene High Bind Microplate (Corning; Cat: 3361) were coated overnight (>16 h) with RH5.1 at 1 μg/mL at 4 °C. Plates were washed with PBST and blocked with 200 μL/well PBS with 2% blotto, non-fat dry milk (PBSM) (Santa Cruz Biotechnology; Cat: sc-2324) for 2 h. Plates were washed and antibodies added at 6 μg/mL diluted in PBSM. Following a 2 h incubation, plates were washed and competing antibodies were added, diluted in PBSM. Competing antibody concentrations were 1.578 nM and 0.162 nM for pR3 and pR5, respectively (set at 2x their *K*_D_ (M) values, as determined by HT-SPR). Following a 2 h incubation, plates were washed and a 1:5000 dilution of goat anti-human IgG (Fab-chain specific) peroxidase conjugate antibody (Millipore Sigma; Cat: A0293) was added and incubated for 1 h. Plates were washed in washing buffer and 50 μL 1-Step™ Ultra TMB-Blotting Solution development buffer (Thermo Scientific; Cat: 37574) was added per well and developed according to internal controls. All mAbs were tested in duplicate. Development was stopped by adding 50 μL 2 M sulfuric acid. Unless otherwise stated, all steps were carried out at r.t.

### Antibody kinetics

High-throughput surface plasmon resonance (SPR) binding experiments were performed on a Carterra LSA instrument equipped with a 2D planar carboxymethyldextran surface (CMDP) chip type (Carterra) using a 384-ligand array format. The CMDP or HC30M chip was first conditioned with 60 s injections of 50 mM NaOH, 1 M NaCl and 10 mM glycine, pH 2.0 before activation with a freshly prepared 1:1:1 mixture of 100 mM 2-Morpholinoethanesulphonic acid (MES), pH 5.5, 100 mM sulfo-N-hydroxysuccinimide, and 400 mM 1-ethyl-3-(3-dimethylaminopropyl) carbodiimide hydrochloride. A coupled lawn of goat anti-human IgG Fc (hFc; 50 μg/mL in 10 mM sodium acetate, pH 4.5) (Jackson ImmunoResearch) was then prepared before quenching with 0.5 M ethanolamine, pH 8.5 and washing with 10 mM glycine, pH 2.0. mAbs prepared at 100 ng/mL in Tris-buffered saline with 0.01 % Tween-20 (TBST) were then captured onto the surface in a 384-array format via a multi-channel device, capturing 96 ligands at a time. For binding kinetics and affinity measurements, an eight-point three-fold dilution series of RH5.1 protein ending at 100 nM in TBST was sequentially injected onto the chip from lowest to highest concentration. For each concentration, the antigen was injected for 5 min (association phase), followed by TBST injection for 15 min (dissociation phase). Two regeneration cycles of 30 s were performed between each dilution series by injecting 10 mM glycine, pH 2.0 on the chip surface. The SPR results were exported to Kinetics Software (Carterra) and analyzed as nonregenerative kinetics data to calculate association rate constant (*k*_on_), dissociation rate constant (*k*_off_) and equilibrium dissociation constant (*K*_D_) values via fitting to the Langmuir 1:1 model. Prior to fitting, the data were referenced to the anti-hFc surface then double referenced using the final stabilizing blank injection.

### Epitope binning by SPR

High-throughput epitope binning experiments were performed in a classical sandwich assay format using the Carterra LSA and an HC30M chip. The chip was conditioned as described above before antibodies prepared at 10 μg/mL in 10 mM sodium acetate, pH 4.5 with 0.05 % Tween were coupled to the surface: the chip surface was first activated with a freshly prepared 1:1:1 activation mix of 100 mM MES, pH 5.5, 100 mM sulfo-N-hydroxysuccinimide, and 400 mM 1-ethyl-3-(3-dimethylaminopropyl) carbodiimide hydrochloride, and antibodies were injected and immobilized onto the chip surface by direct coupling. The chip surface was then quenched with 1 M ethanolamine, pH 8.5, followed by washing with 10 mM glycine, pH 2.0. Sequential injections of 50 nM RH5.1 protein (5 min) followed immediately by the 10 μg/mL sandwiching antibody (5 min), both diluted in TBS-T, were added to the coupled array and the surfaces regenerated with 10 mM glycine, pH 2.0 using two 30 s regeneration cycles. Using the Carterra Epitope software, normalized response unit (RU) of mAb pairs were automatically sorted into a heatmap readout, which were further used to plot a dendrogram with a cut-off height set to cluster antibodies into monophyletic communities.

### Growth inhibition activity (GIA) assay

Single concentration *in vitro* GIA assays were carried as previously described according to the methods of the GIA International Reference Centre at NIAID/NIH, USA.^11^ All assays used 3D7 clone *P. falciparum* parasites cultured in human RBC from in-house volunteer donations or supplied by the UK NHS Blood and Transplant service for non-clinical issue. Briefly, mAbs were buffer exchanged into incomplete parasite growth media (RPMI, 2 mM L-glutamine, 0.05 g/L hypoxanthine, 5.94 g/L HEPES) before performing the GIA assay and allowing parasites to go through a single cycle of growth. To ensure consistency between experiments, in each case the activity of a negative control human mAb, EBL040,^63^ which binds to the Ebola virus glycoprotein, and three anti-PfRH5 mAbs with well-characterized GIA (2AC7, R5.016, and R5.034)^29,30^ were run alongside the test mAbs, and were all tested in triplicates. mAbs showing >30 % GIA were subsequently tested in an eight step, five-fold dilution series with a final assay start concentration of 1 mg/mL to determine interpolated EC values. The resultant data were transformed according to x=log(x) and the transformed data were fitted by an asymmetrical sigmoidal five-parameter non-linear regression. GIA values were interpolated from the resultant curve. If a mAb did not reach a sufficiently high GIA (i.e., the mAb did not reach 50 % or 80 % at any test concentration), then it was assigned a “negative” value of 10 mg/mL for that particular EC readout.

For the OREs, mAbs were pooled at relative concentrations (normalized to each other) as determined by Ig-Seq, as described in Data S1. When a mAb was removed from a pool condition, it was substituted with negative control mAb EBL040^63^ to keep the concentrations of anti-RH5.1 mAbs constant. All pools were tested in an eight step, two-fold dilution series with a final assay start concentration of 0.5 mg/mL to determine interpolated EC values. As done with individual mAbs, the resultant data were fitted and GIA values were interpolated from the resultant curves, with insufficient GIA readout, or “negative” GIA, valued at 10 mg/mL. OREs were run across triplicate experiments, each with a distinct donor blood-batch, to minimize noise of the assay.^64^ EC values were averaged across all three experiments and standard deviation was calculated.

For synergy screens of intra-PfRH5 mAb interactions with pR5.003 and pR5.005 shown in Fig. 3, single concentration assays were carried out as above, with neutralizing antibodies added at a concentration equivalent to their interpolated EC_50_ value and non-neutralizing antibodies held at 0.5 mg/mL. pR5.003 and pR5.005 were held constant at 0.2 mg/mL individually or 0.4 mg/mL when added together (each 0.2 mg/mL). For synergy screens of intra-PfRH5 mAb interactions, pR3 and/or pR5 were held at either their ORE concentrations (0.06 and 0.05 mg/mL, respectively) or at 0.3 mg/mL. Test (pR1, pR, pR4) and control mAbs (pR3 and pR5) were held at their respective ORE concentrations. When tested together, pR3 and pR5 were held individually at 0.25 mg/mL, for a final concentration of 0.5 mg/mL.

For analysis of synergistic or antagonistic interactions, the Bliss additivity^65,66^ was determined based on the measured activity from each antibody alone (1 and 2) using the following formula:

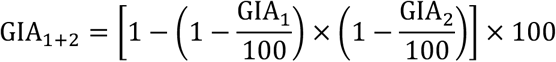

### Statistical Analysis

Analysis was performed using GraphPad Prism version 10.2.1 (GraphPad Software, LLC). Tests and statistics are described in figure legends. Non-parametric tests were chosen for non-normally distributed data. In all statistical tests, reported P values are two-tailed and P < 0.05 considered significant.

## ACKNOWLEDGEMENTS

The authors thank Kate Skinner, Sarah Silk, Jenny Bryant and Lana Strmecki (University of Oxford); Lorraine Soisson and Robin Miller (USAID); Rick King, Tara Tagmyer and Ashley Birkett (PATH); and the VAC063 trial participants.

This work was funded in part by the United States Agency for International Development (USAID) Malaria Vaccine Development Program (MVDP) (7200AA20C00017), the findings and conclusions are those of the authors and do not necessarily represent the official position of USAID; the UK Medical Research Council (MRC) [MR/X012085/1]; the National Institute for Health Research (NIHR) Oxford Biomedical Research Centre (BRC) and NHS Blood & Transplant (NHSBT; who provided material), the views expressed are those of the authors and not necessarily those of the NIHR or the Department of Health and Social Care or NHSBT.

## AUTHOR CONTRIBUTIONS

Conceived and performed experiments and/or analyzed the data: JM, KM, JRB, AS, CR, DQ, AR, AH, DP, JK, DRT, SAK.

Project management: RSM.

Contributed reagents, materials, and analysis tools: AMM.

Wrote the paper: JM, KM, GG, GCI, SJD, JJL.

